# Matrix viscoelasticity regulates dendritic cell migration and immune priming

**DOI:** 10.1101/2025.09.28.678961

**Authors:** Wei-Hung Jung, Emie Humann, Joshua M Price, Yoav Binenbaum, Azra Haseki, Sanjana Iyer, Daivd J Mooney

**Affiliations:** John A. Paulson School of Engineering and Applied Sciences, Harvard University, Cambridge, MA 02138, USA; Wyss Institute for Biologically Inspired Engineering, Harvard University, Boston, MA 02215, USA; Dana-Farber/Boston Children’s Cancer and Blood Disorders Center, Boston, MA 02215, USA

## Abstract

The tumor microenvironment shapes immune surveillance through its mechanical properties, yet the role of matrix viscoelasticity remains unclear. Here, we used a collagen system with tunable viscoelasticity to define how matrix relaxation directs dendritic cell (DC) behavior. Elastic matrices impaired DC migration by limiting actomyosin-driven collagen remodeling, thereby reducing DC-T cell encounters and weakening T cell priming, activation, proliferation, and tumor killing. Blocking DC migration in fast-relaxing gels recapitulated key aspects of the impaired T cell priming seen in elastic matrices. Prolonged confinement in elastic extracellular matrix induced a mechanomemory state, locking DCs into reduced motility even after transfer to viscoelastic environments, corresponding to altered chromatin accessibility. Finally, studies with patient-derived ependymoma samples confirmed these findings, identifying viscoelasticity as a barrier to antitumor immunity with implications for therapeutic intervention.

## Introduction

Despite the presence of tumor-infiltrating lymphocytes, T cells often fail to control tumor growth and metastasis^1-3^. This failure reflects not only tumor-intrinsic resistance mechanisms but also inadequate priming and support from the tumor microenvironment (TME)^4,5^. Dendritic cells (DCs) are central regulators of antitumor immunity, bridging innate recognition with adaptive T cell responses^6,7^. In tumors and fibrotic tissues, however, DCs frequently exhibit impaired function, limiting their ability to license productive T cell responses^8-10^. Effective T cell priming and activation require that DCs migrate through dense stroma, remodel extracellular matrix (ECM), and establish contacts with T cells^11-16^. When these processes are compromised, T cells cannot mount robust effector responses, contributing to the poor immune surveillance characteristic of many solid tumors.

The TME is a complex milieu where biochemical and physical cues converge to shape immune responses^17-21^. Physical properties of the ECM, including stiffness, architecture, and viscoelasticity, have emerged as potent regulators of cell behaviors^22-28^. Among these, viscoelasticity, a time-dependent response to deformation, remains comparatively underexplored. Perturbations in viscoelasticity occur in fibrotic disease and cancer, where excessive collagen deposition and crosslinking increase matrix elasticity and prolong stress relaxation^29-34^, features associated with reduced immune infiltration and an immunologically “cold” phenotype^35,36^. Mechanical cues can also leave lasting effects, or “mechanomemory,” through epigenetic remodeling^37-39^. Recent studies have shown that viscoelasticity can contribute to mechanomemory by enhancing plasticity in fibroblasts^40^, or by stably imprinting gene expression programs in T cells^41^. Whether DCs, whose migration and priming of T cells are uniquely essential for antitumor immunity, are subject to similar regulation remains unknown. This represents a critical gap, as mechanical regulation of DCs could directly constrain their ability to initiate effective immune responses.

In this study, we hypothesized that altered viscoelasticity shapes DC function and limits T cell priming, contributing to impaired antitumor immunity. Using a type I collagen system, the principal ECM component of tumor stroma and fibrotic tissue^42,43^, in which viscoelasticity can be tuned independently of stiffness, we found that DC migration was impaired in slow-relaxing matrices owing to reduced actomyosin-driven collagen remodeling. This confinement curtailed DC-T cell interactions, reducing T cell proliferation, activation, and tumor cell killing. Prolonged confinement in slow-relaxing gels induced a form of mechanomemory, whereby DCs retained restricted migration even after transfer to permissive environments, linked to reduced chromatin accessibility at mechanosensing loci. Inhibiting DC migration fast-relaxing gels reduced T cell activation, recapitulating the impaired priming observed in slow-relaxing gels and underscoring migration as critical for T cell priming. Finally, in patient-derived carcinoma samples, viscoelastic regulation similarly constrained DC-mediated T cell activation, highlighting viscoelasticity as a barrier to antitumor immunity and a target to improve DC-based immunotherapies.

## Results

### ECM elasticity controls DC migration and gene expression

To investigate how ECM viscoelasticity regulates DC behavior, type I collagen hydrogels were fabricated in which stress relaxation can be tuned independently of stiffness using bio-orthogonal click chemistry. Collagen was functionalized with norbornene, enabling site-specific crosslinking through an inverse electron-demand Diels– Alder reaction with a methyltetrazine-(PEG)_5_-methyltetrazine (divalent MeTz) crosslinker (**Fig.1a**)^41^. Confocal imaging revealed an average pore size of 3.1 μm, which was unchanged between non-crosslinked (Fast R) and crosslinked (Slow R) gels (**Fig.1b**). Rheological measurements showed that crosslinking extended the stress relaxation time of 2 mg mL^−1^ collagen gels from 72 to 1,252 s without altering the storage modulus (**Fig.1c,d**). The loss angle decreased in crosslinked gels, indicating a more elastic network (**Fig.1e**). A similar extension of relaxation time without changes in storage modulus was also observed in gels with higher collagen concentration (4 mg mL^−1^) (**Supplementary Fig.1a-c**). Thus, this system permits independent control of viscoelasticity while preserving stiffness and microarchitecture. We next asked whether viscoelasticity influences DC migration. Human monocyte-derived DCs were differentiated from monocytes isolated from peripheral blood mononuclear cells (PBMCs), activated with TNF-α, and encapsulated in 2 mg mL^−1^ Fast R or Slow R gels (**Fig.1f,g**). Time-lapse imaging showed that DCs in Slow R gels migrated with confined trajectories compared with cells in Fast R gels. Mean squared displacement (MSD) analysis showed a marked decrease in cell displacement in Slow R gels, compared to Fast R gels (**Fig.1h**). Similarly, DCs encapsulated in higher-concentration gels (4 mg mL^−1^) demonstrated restricted migration in the Slow R condition (**Supplementary Fig.1d**,**e**).

**Fig. 1.**
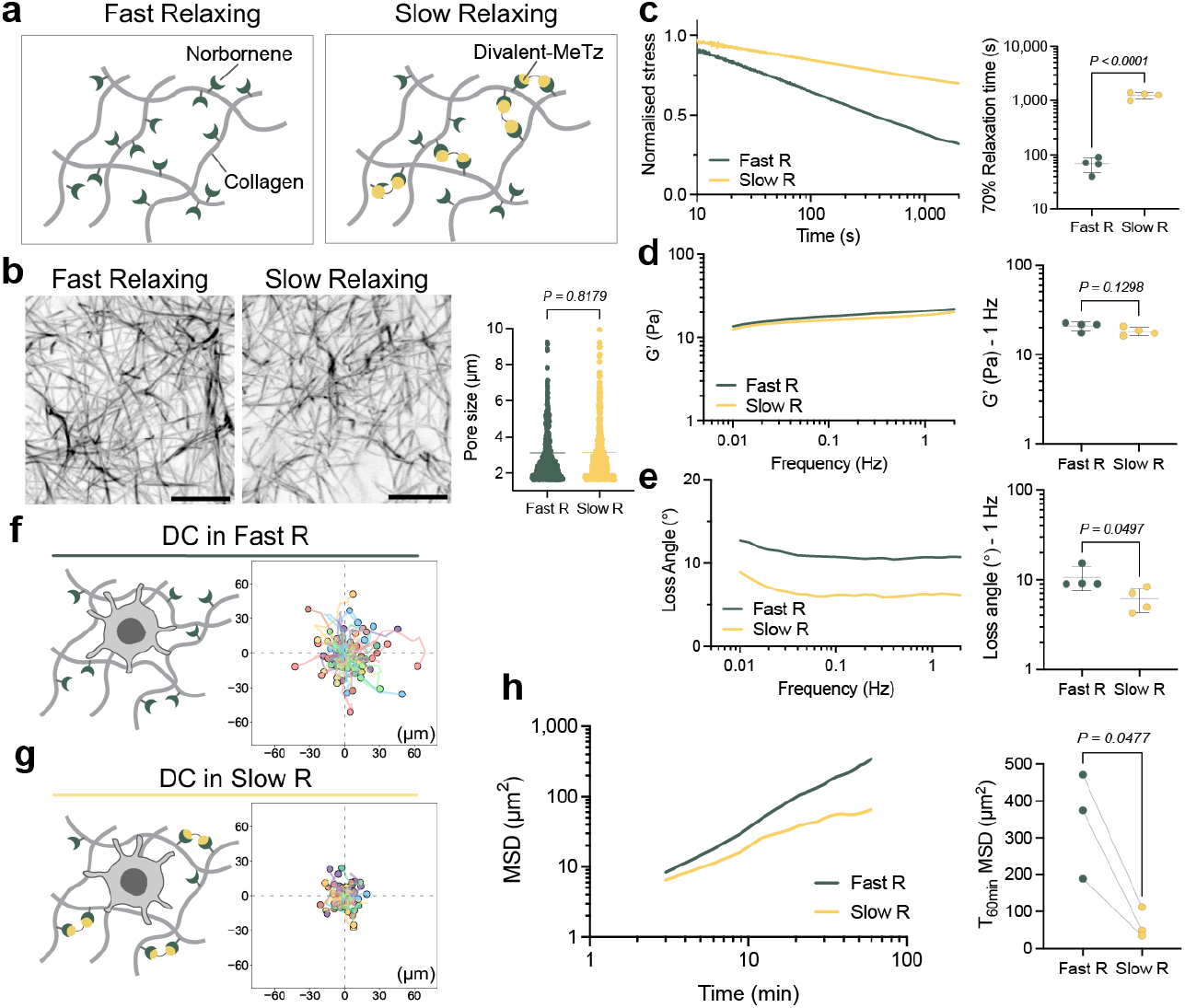
Human DC migration is confined in slow-relaxing collagen gels. **a**, Schematic illustrating the structure of fast-relaxing norbornene-functionalized 2 mg mL^−1^ collagen gels (Fast R) and localized crosslinking with a divalent MeTz crosslinker to generate slow-relaxing gels (Slow R). **b**, Representative confocal microscopy images of Fast R and Slow R collagen gels labeled with tetrazine-iFluor 488, and quantification of gel pore size from 3 independently formed gels; lines indicate the mean. *P* value was determined using a two-tailed Mann–Whitney test. Scale bar: 10 μm. **c**, Stress relaxation profiles of 2 mg mL^−1^ Fast R and Slow R gels, and quantification of the time required for stress to decay to 70% of the initial value. **d**, Storage modulus (G’) of Fast R and Slow R gels determined by frequency sweep, and corresponding quantification at 1 Hz. **e**, Loss angle of Fast R and Slow R gels determined by frequency sweep, with quantification at 1 Hz. Results in **c-e** are from four independently formed gels and are presented as mean ± s.d. *P* values were determined using a two-tailed unpaired t-test. **f, g**, Schematic of DCs encapsulated in Fast R or Slow R gels, and representative migration trajectories over 60 minutes. **h**, Quantification of DC MSD over time within Fast R or Slow R gels, with differences in MSD values highlighted at the 60-minute time point. Data are representative of three independent donors; lines connect values from the same donor. *P* values were determined using a two-tailed paired t-test (*n* = 3).

To examine how ECM viscoelasticity shapes DC transcriptional programs, activated DCs were encapsulated in 2 mg mL^−1^ Fast R or Slow R hydrogels for 3 days, followed by bulk RNA sequencing (**Fig.2a**). Principal component analysis of gene expression in four donors showed that ECM viscoelasticity was the main factor separating transcriptomes, with PC1 distinguishing relaxation conditions and PC2 capturing inter-donor variability (**Fig.2b**). Gene ontology analysis of genes differentially expressed between Fast R and Slow R DCs revealed enrichment of biological processes related to cytokine production, T cell proliferation, epithelial-to-mesenchymal transition, and extracellular structure organization (**Fig.2c**). Heatmap analysis of cytokine production–related genes across donors revealed higher expression of Toll-like receptors 7 (TLR7) in Fast R, whereas TLR1 and TLR4 were more highly expressed in Slow R (**Fig.2d**). Transcription factor activity analysis showed RE1-silencing transcription factor, a regulator of chromatin remodeling^44^, as the most activated regulator in Fast R DCs, whereas AP-1 (JUN, FOS, JUNB), C/EBP (CEBPB, CEBPD), and NF-κB (RELA, NFKB1) were among the top predicted regulators in Slow R DCs (**Fig.2e**). Differential expression analysis showed significant upregulation of key AP-1 and NF-κB target genes, including IL1-β and IL-6, two pro-inflammatory cytokines co-regulated by both pathways (**Fig.2f**). To validate these findings at the protein level, activated DCs were encapsulated in fast- or slow-relaxing gels for 3 days and culture supernatants collected for cytokine detection. Consistent with transcriptomic data, both IL-1β and IL-6 secretion were significantly elevated in the slow-relaxing condition (**Fig.2g**).

**Fig. 2.**
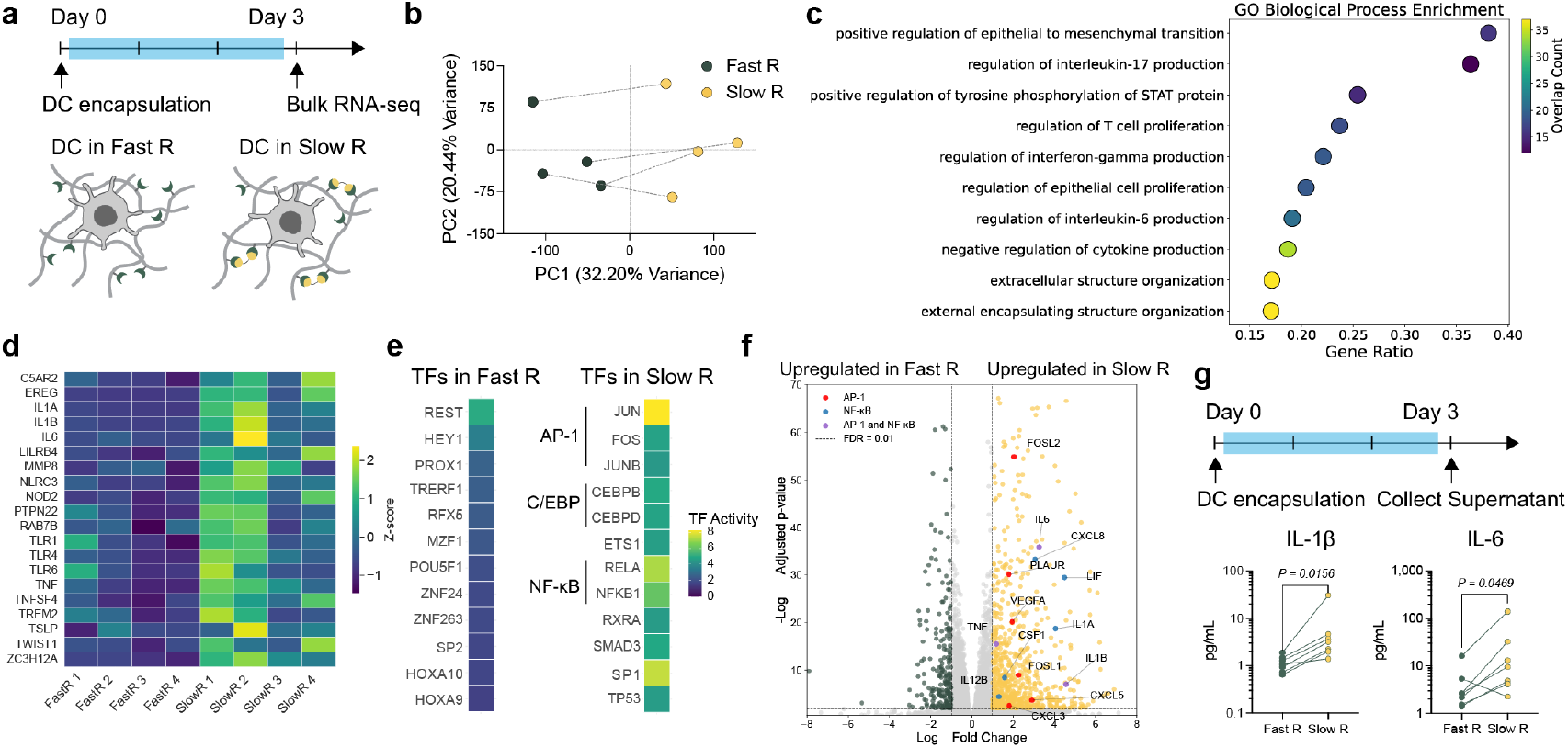
Viscoelastic properties of the ECM differentially regulate DC gene expression and transcription factor activation. **a**, Schematic of the experimental timeline for bulk RNA-seq analysis. DCs were encapsulated in 2 mg mL^−1^ Fast R or Slow R hydrogels for 3 days, followed by collagen gel digestion and RNA extraction. **b**, Principal component analysis of bulk RNA-seq data from four independent donors. Lines connect paired Fast R and Slow R samples from the same donor. **c**, GO biological process enrichment analysis of the top 10 pathways derived from genes differentially expressed between DCs in Fast R and Slow R hydrogels. **d**, Heatmap of genes associated with the “regulation of interleukin-6 production” GO term. Numbers indicate donor identity. **e**, Transcription factor activity analysis of the top 12 transcription factors upregulated in Fast R and Slow R conditions. Transcription factors belonging to the AP-1, C/EBP, and NF-κB families are highlighted. **f**, Volcano plot of differentially expressed genes between Fast R and Slow R conditions. Genes upregulated in the Slow R condition and associated with AP-1 (red), NF-κB (blue), or both (purple) pathways are highlighted. **g**, Schematic of the experimental timeline for supernatant collection and ELISA measurement of IL-1β and IL-6. Data are from seven independent donors. Lines connect paired Fast R and Slow R samples from the same donor. *P* values were determined using a two-tailed paired t-test (*n* = 7).

### Reduced T cell priming ability of DCs in elastic ECM

Given the central role of DCs in priming T cells, we next asked whether ECM viscoelasticity modulates the ability of DCs to initiate T cell responses. DCs were pulsed with SK-MEL-5 melanoma lysates and subsequently activated with TNF-α. Autologous human pan-T cells were then co-encapsulated with DCs in either Fast R or Slow R gels and cultured for 12 days (**Fig.3a**). T cell proliferation was greater in Fast R gels, with both CD4^+^ and CD8^+^ subsets expanding more robustly than in Slow R gels (**Fig.3b,c**). A higher fraction of CD4^+^ T cells expressed the activation markers CD25 and OX40 in Fast R gels compared with those primed in Slow R gels (**Fig.3c,d**). Similarly, a greater proportion of CD8^+^ T cells expressed CD25 and PD-1 in Fast R gels (**Fig.3e,f**), indicating that both CD4^+^ and CD8^+^ subsets were less effectively primed by DCs in slow-relaxing ECM. To test whether these differences reflected a direct effect of ECM viscoelasticity on T cells, we encapsulated T cells alone in Fast R or Slow R gels (**Supplementary Fig.2a**). CD4^+^ and CD8^+^ T cell proliferation was comparable across these two conditions, and CD25 expression was also similar and lower than in the presence of DCs (**Supplementary Fig.2b**,**c**). These findings indicate that the effects of viscoelasticity on T cell activation here are mainly mediated through DCs.

**Fig. 3.**
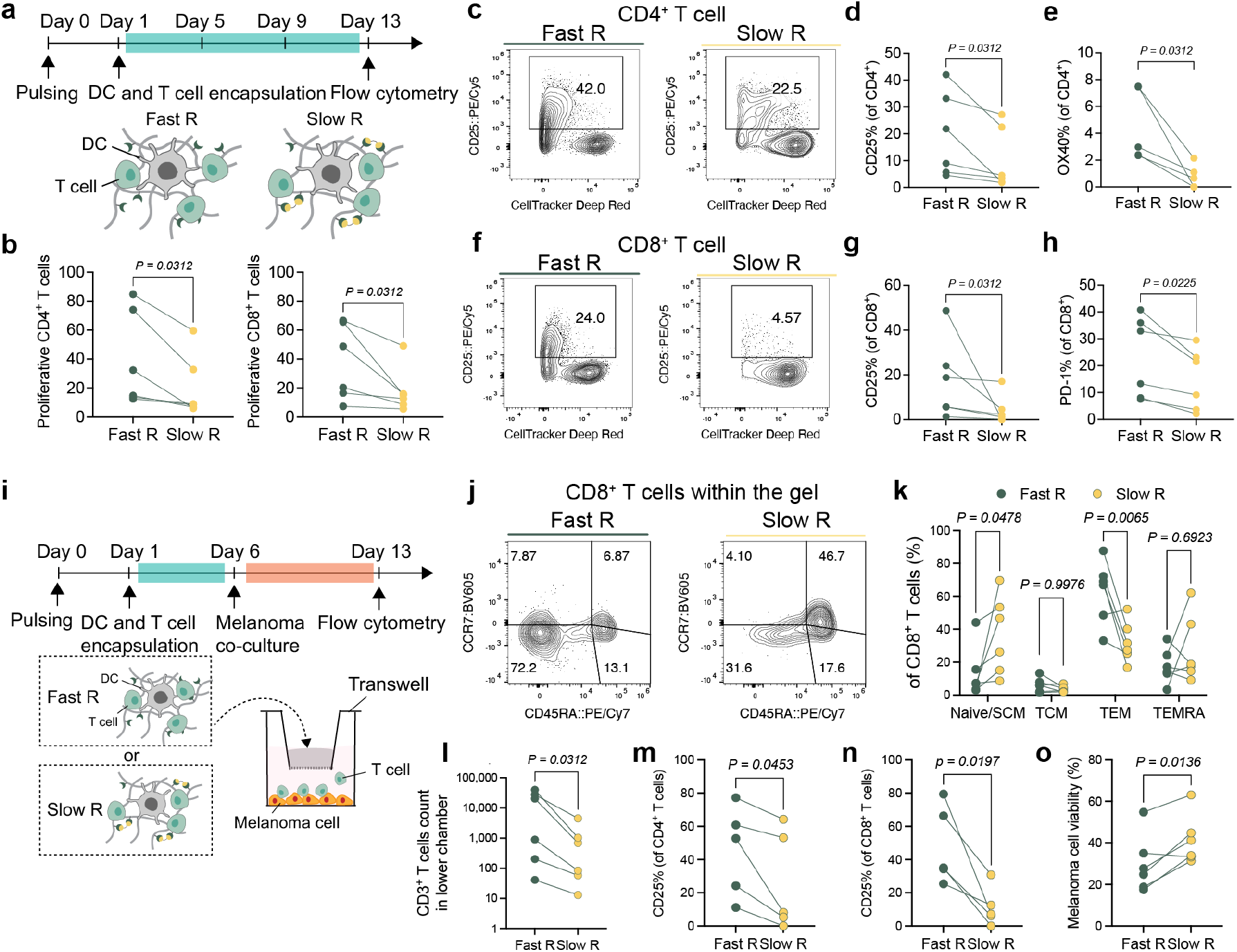
ECM viscoelasticity impairs DC-mediated T cell priming, reduces T cell activation, and limits cancer cell killing. **a**, Schematic and timeline of DCs and initially unactivated T cells encapsulated in 2 mg mL^−1^ Fast R or Slow R hydrogels. At day 0, DCs were pulsed with SK-MEL5 melanoma lysates and activated with TNF-α. At day 1, DCs were co-encapsulated with unactivated T cells for an additional 12 days. **b**, Quantification of proliferating CD4^+^ and CD8^+^ T cells, measured as reduction in deep red dye signal after 12 days of co-culture. *P* values were determined using a two-tailed Wilcoxon matched-pairs signed rank test (*n* = 6). **c**, Representative flow cytometry plots showing greater CD25^+^ CD4^+^ T cells and proliferation (deep red signal) in Fast R. **d**,**e**, Quantification of CD25^+^ or OX40^+^ CD4^+^ T cells, as measures of activation. *P* values were determined using a two-tailed Wilcoxon matched-pairs signed rank test (*n* = 6). **f**, Representative flow cytometry plots showing increased CD25^+^ CD8^+^ T cells and proliferation (deep red signal) in Fast R. **g**,**h**, Quantification of CD25^+^ or PD-1^+^ CD8^+^ T cells as measures of activation. *P* values were determined using a two-tailed Wilcoxon matched-pairs signed rank test for CD25 and a two-tailed paired t-test for PD-1 (*n* = 6). **i**, Schematic and timeline of DCs co-encapsulated with initially unactivated T cells in Fast R or Slow R matrices for 12 days in a transwell assay; at day 6, inserts were transferred to wells containing SK-MEL5 melanoma cells. **j**, Representative flow cytometry plots of CD8^+^ T cell subsets after 12 days in Fast R and Slow R hydrogels. **k**, Quantification of CD8^+^ T cell subsets within the gel after 12 days of culture, defined by CD45RA and CCR7 expression (Naïve/SCM: CCR7^+^CD45RA^+^; TCM: CCR7^+^CD45RA^−^; TEM: CCR7^−^CD45RA^−^; TEMRA: CCR7^−^CD45RA^+^). *P* values were determined using a two-way repeated-measures ANOVA (*n* = 6 donors, each measured across all conditions), followed by Šídák’s multiple comparisons test to assess differences between Fast R and Slow R for each T cell subset. **l**, Quantification of CD3^+^ T cells that migrated to the bottom chamber after 7 days of co-culture with melanoma cells. **m**,**n**, Quantification of CD4^+^ and CD8^+^ T cells expressing CD25 following migration through the transwell. **o**, Viability of melanoma cells after 7 days of co-culture with T cells. Lines connect paired data points from the same donor in the Fast R and Slow R groups. The *P* value for **l** was determined using a two-tailed Wilcoxon matched-pairs signed rank test, and *P* values for **m–o** were determined using a two-tailed paired t-test (*n* = 6 for **l**,**o**; *n* = 5 for **m**,**n**).

To further evaluate T cell function and migratory capacity, DC-T cell encapsulated co-cultures were placed in transwells (4 μm pore size). After 5 days, inserts were transferred to wells containing SK-MEL-5 cells and cultured for an additional 7 days, allowing activated T cells to migrate and engage tumor cells (**Fig.3i**). Flow cytometric analysis showed that CD8^+^ T cells retained within Slow R gels exhibited a greater naïve phenotype (CD45RA^+^CCR7^+^) and reduced effector memory phenotype (CD45RA^−^CCR7^−^), as compared to those in Fast R gels (**Fig.3j,k**). Fewer T cells migrated from Slow R gels to the tumor cell compartment, as compared to T cells in Fast R gels (**Fig.3l**). Among the migrated T cells in the Slow R condition, significantly fewer CD4^+^ and CD8^+^ T cells were activated (**Fig.3m,n)**, and this correlated with reduced tumor cell killing in this condition (**Fig.3o**). The effect of viscoelasticity on T cells was also evaluated by encapsulating T cells alone in the transwell system. In this setting, all four T cell subsets remained comparable across conditions (**Supplementary Fig.3a**,**b**), and migrated T cell numbers, activation status, and tumor cell viability were also similar (**Supplementary Fig.3c–f**). Together, these findings indicate that a more elastic ECM diminishes the ability of DCs to activate T cells, thereby limiting T cell migration and cytotoxicity.

To examine whether diminished DC-T cell contacts underlie the reduced T cell priming in Slow R, SK-MEL-5-pulsed DCs and autologous T cells were co-encapsulated in Fast R or Slow R gels and imaged by time-lapse microscopy (**Supplementary Fig.4a**). DC-T cell interactions (co-localization) were consistently more frequent in Fast R gels (**Supplementary Fig.4b**). Consistent with earlier findings, DCs displayed significantly reduced motility in Slow R gels (**Supplementary Fig.4d–f**), whereas T cell migration was unaffected when co-cultured with DCs (**Supplementary Fig.4g-i**) or encapsulated alone (**Supplementary Fig.5**). To test whether impaired priming reflected altered DC activation, we examined co-stimulatory and antigen-presenting molecule expression after 3 days of encapsulation and observed no substantial differences between conditions (**Supplementary Fig. 6**). Together, these results suggest that reduced DC motility in slow-relaxing ECM limits the frequency of DC-T cell encounters, thereby impairing effective T cell priming.

### ECM viscoelasticity induces lasting changes in DC migration and chromatin accessibility

We next investigated whether ECM viscoelasticity leads to persistent changes in DC migration. DCs were conditioned in Fast R or Slow R gels for 1 or 3 days, then re-encapsulated in Fast R gels (**Fig.4a**). DCs conditioned for 1 day showed no difference in migration after transfer (**Fig.4b,c)**. In contrast, DCs pre-exposed to Slow R for 3 days exhibited reduced motility that persisted for up to 3 days post-transfer (**Fig.4d,e**). To probe the basis for these changes, DCs were again encapsulated in Fast R or Slow R gels for 3 days, followed by ATAC-seq analysis (**Fig.4f**). Global accessibility around transcription start sites (TSSs) was comparable between conditions (**Supplementary Fig.7**). However, analysis revealed reduced accessibility of mechanosensing-associated genes, (**Fig.4g**) including WWTR1 (TAZ) and WASF1, a positive regulator of Arp2/3-mediated actin nucleation^45^, in Slow R gels (**Fig.4h**). These results suggest that prolonged exposure to a slow-relaxing ECM induces stable changes in chromatin accessibility that impair DC mechanosensing and migration, even after removal from the original environment.

**Fig. 4.**
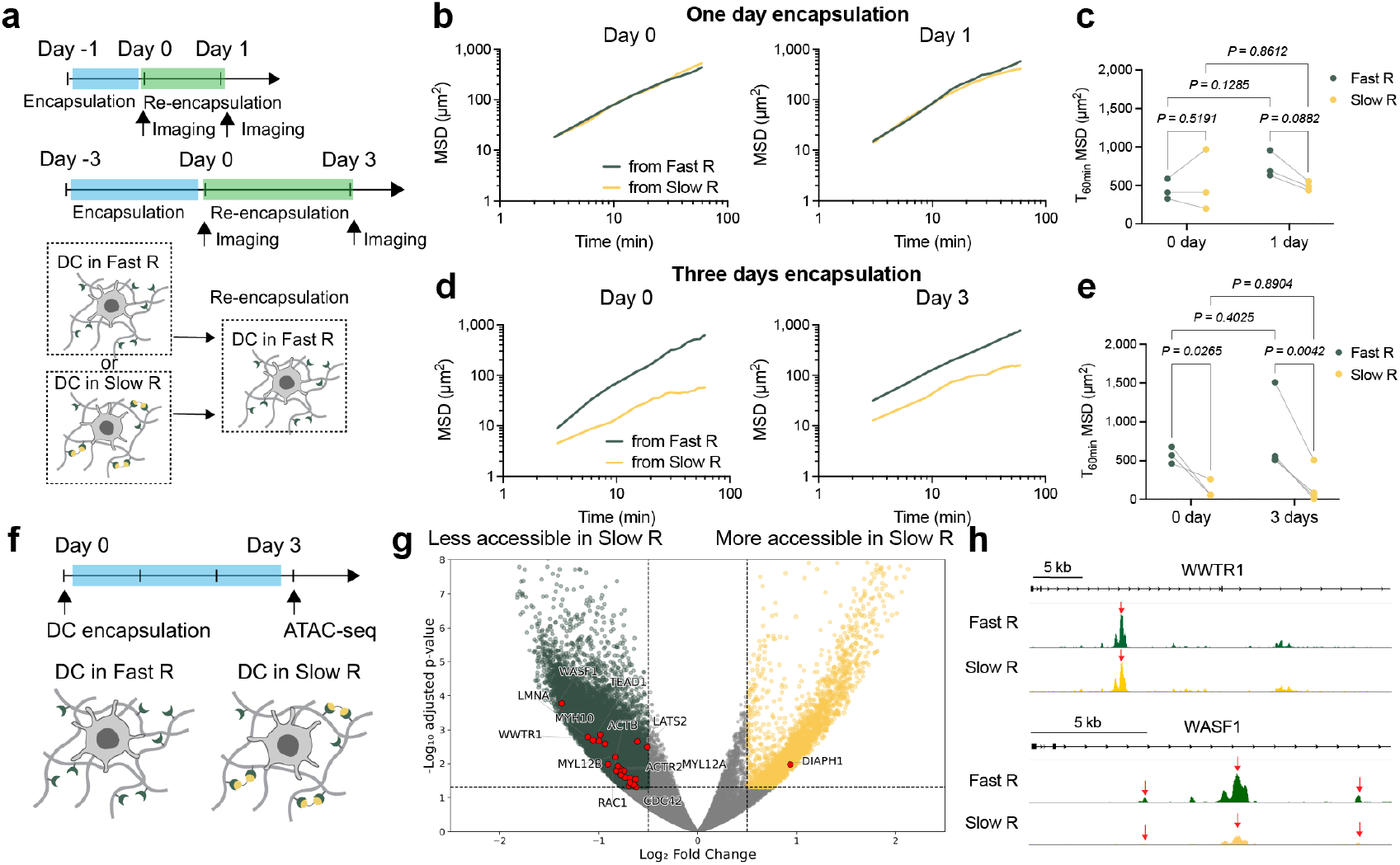
ECM elasticity induces lasting changes in DC migration and mechanosensing pathway accessibility. **a**, Schematic of the experimental timeline for re-encapsulation experiments. DCs were first conditioned in Fast R or Slow R for 1 or 3 days, followed by collagen digestion and re-encapsulation in Fast R. For 1-day conditioning, imaging to evaluate DC migration was performed after 2 hours or 1 day post-re-encapsulation. For 3-day conditioning, imaging was performed after 2 h or 3 days post-re-encapsulation. **b-e**, Quantification of MSD over time for DCs conditioned in Fast R or Slow R for 1 day (**b**,**c**) or 3 days (**d**,**e**) and re-encapsulated in Fast R, with MSD values at 60 min shown. Data are from three independent donors; lines connect paired values from the same donor. *P* values were determined using a two-way repeated-measures ANOVA with uncorrected Fisher’s LSD (*n* = 3 donors, each measured across all conditions). **f**, Schematic of the experimental timeline for ATAC-seq analysis. DCs were encapsulated in 2 mg mL^−1^ Fast R or Slow R hydrogels for 3 days, followed by collagen digestion and library preparation for sequencing. **g**, Volcano plot of differentially accessible regions between Fast R and Slow R conditions. Genes related to mechanosensing pathways that are highlighted in red. **h**, Representative gene accessibility tracks for WWTR1 and WASF1. Red arrows indicate reduced accessibility in Slow R. Boxes above the gene accessibility denote promoter, exon, and intron regions; arrows indicate the direction of transcription. Coverage tracks are shown with the y-axis scaled from 0 to 500 reads.

### DC migration in ECM requires actomyosin contractility and collagen remodeling

Given the reduced chromatin accessibility of actomyosin-related genes identified by ATAC-seq, we next examined the functional role of actomyosin contractility and matrix remodeling in DC migration. DCs were first treated with para-amino-blebbistatin (a myosin II inhibitor) or Y-27632 (a ROCK inhibitor). Both treatments significantly reduced MSD in Fast R gels but had little effect in Slow R gels (**Fig.5a-c**). Similarly, inhibiting focal adhesion kinase with defactinib or actin polymerization with latrunculin A markedly impaired DC migration in Fast R gels (**Fig.5d-f**), supporting the importance of actomyosin contractility for migration in permissive viscoelastic environments. However, inhibition of PI3K, previously linked to mechanosensing in monocyte-to-DC differentiation^46^, did not affect DC migration (**Supplementary Fig.9**), further supporting a dominant role for actomyosin contractility.

We next probed DC interactions with the surrounding ECM. To determine whether DC-mediated matrix degradation was required for migration in viscoelastic gels, cells were treated with broad-spectrum MMP inhibitors (GM6001, marimastat). However, these had no effect on MSD (**Supplementary Fig.8**), indicating that early DC migration is largely MMP-independent. The ability of DCs to deform the surrounding collagen were then tracked by time-lapse confocal imaging. In Fast R gels, DCs generated robust collagen displacements that were largely absent in Slow R gels (**Fig. 5g**,**h; Supplementary Movies1**,**2**). Inhibition of myosin II with para-amino-blebbistatin abolished collagen remodeling in Fast R gels, phenocopying the behavior of DCs in Slow R matrices (**Fig.5i,j; Supplementary Movies3**,**4**). Quantitative particle image velocimetry revealed extensive directional collagen remodeling extending >35 µm from the cell body in Fast R gels, whereas remodeling was minimal in Slow R gels or under para-amino-blebbistatin treatment (**Fig.5k,l**). Together, these results show that DC migration in fast-relaxing ECM is driven by actomyosin-dependent contractility and requires active collagen remodeling, while in slow-relaxing gels reduced matrix deformability likely constrains collagen remodeling and limits DC motility.

**Fig. 5.**
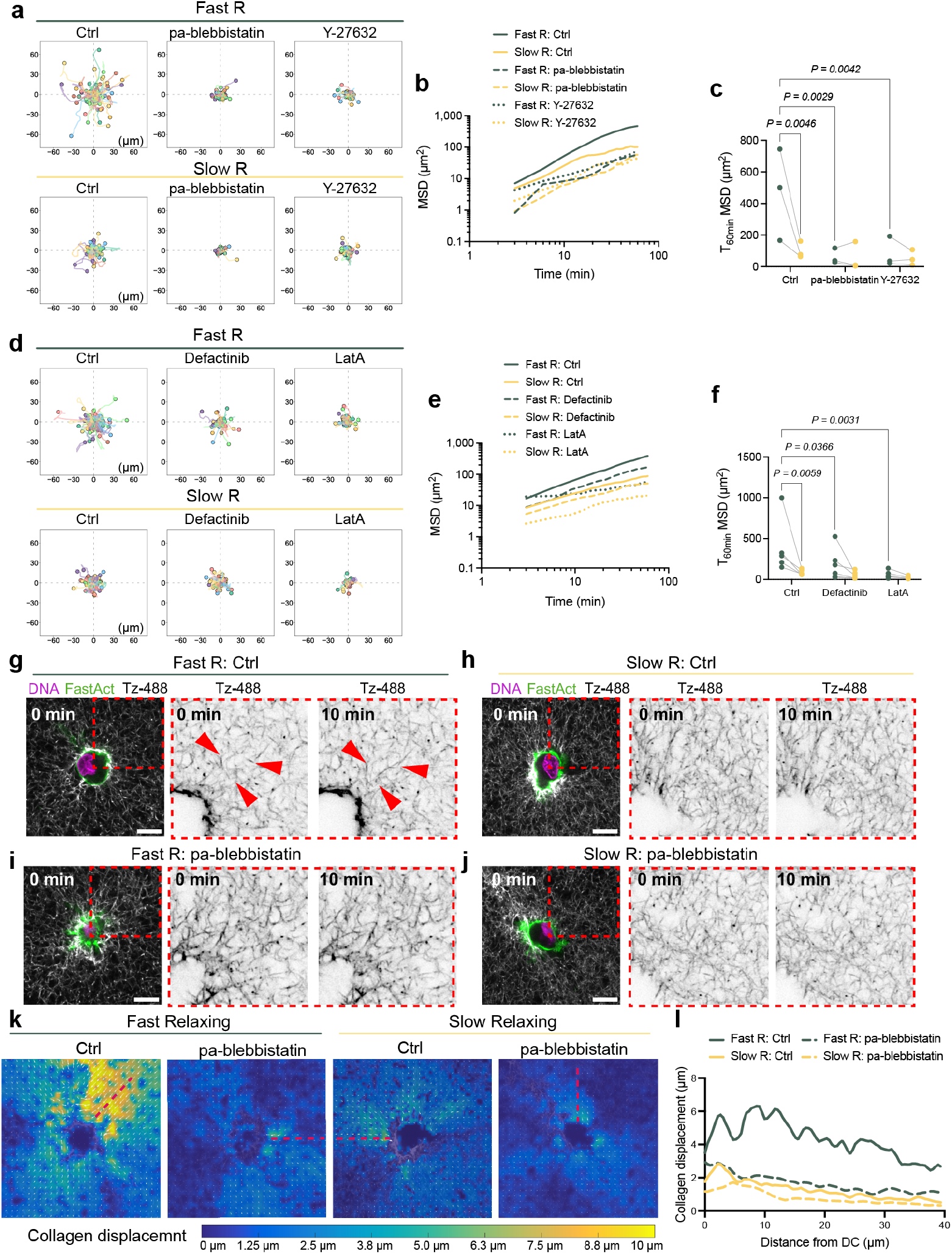
DC migration depends on collagen remodeling through actomyosin contractility. **a**, Representative trajectories of DCs treated with DMSO (Ctrl), pa-blebbistatin, or Y-27632 and encapsulated in 2 mg mL^−1^ Fast R or Slow R gels over 60 min. **b**,**c**, Quantification of DC MSD over time, with MSD values at 60 min shown in paired donor samples; lines connect values from the same donor. **d**, Representative trajectories of DCs treated with DMSO (Ctrl), defactinib, or latrunculin A and encapsulated in 2 mg mL^−1^ Fast R or Slow R gels over 60 min. **e**,**f**, Quantification of DC MSD over time, with MSD values at 60 min shown in paired donor samples; lines connect values from the same donor. *P* values for **c, f** were determined using a two-way repeated-measures ANOVA with uncorrected Fisher’s LSD, with viscoelasticity and treatment as factors, and each donor measured across all conditions (*n* = 3 for **c**; *n* = 5 for **f**). **g–j**, Time-lapse images of DCs encapsulated in 2 mg mL^−1^ Fast R or Slow R gels and treated with DMSO (Ctrl) or para-blebbistatin. Nuclei and actin were labeled with SiR-DNA (magenta) and LifeAct (green), respectively, and collagen fibrils were labeled by clicking Tetrazine-iFluor 488. Collagen remodeling by DCs is shown at time 0 and 10 min, with red arrows indicating prominent remodeling. Scale bar, 10 µm. **k**, Representative PIV plot of DC-mediated collagen displacement over the 10 min imaging period, with the DC located at the center. Displacement is color-coded from 0 μm (blue) to 10 μm (yellow). The dotted red line indicates the direction of most prominent displacement, used for quantification in **l. l**, Quantification of collagen displacement along the direction of most prominent remodeling across the four conditions (*n* = 9–10 per condition).

### Inhibition of DC migration reduces T cell activation

To test whether reduced T cell priming in Slow R gels is directly attributable to impaired DC migration, we engineered DCs to undergo migration arrest using a bio-orthogonal click ligation strategy. DCs were surface-functionalized with MethylTetrazine-PEG_4_ (MeTz) via NHS chemistry, generating MeTzDCs capable of covalently binding to the norbornene-modified collagen in Fast R. The bound MeTz remained largely confined to the cell surface (**Fig.6a,b**). In 2 mg mL^−1^ collagen (no norbornene-modification of the collagen), MeTzDCs migrated comparably to unmodified DCs (**Supplementary Fig. 10**), indicating that MeTz conjugation alone did not alter DC motility. In contrast, encapsulation of MeTzDCs in Fast R resulted in confined trajectories and reduced MSD (**Fig. 6c,d**), consistent with migration arrest in Slow R gel. To directly test whether confining DC migration impairs priming of T cells, MeTzDCs or control DCs were co-cultured with autologous T cells for 3 days, a duration selected based on the retention of MeTz on the DC surface (**Supplementary Fig.11**). Flow cytometry revealed a significant reduction in CD25^+^ CD4^+^ T cells primed by MeTzDCs, demonstrating that limiting DC migration can contribute to impaired T cell activation (**Fig. 6e,f**).

**Fig. 6.**
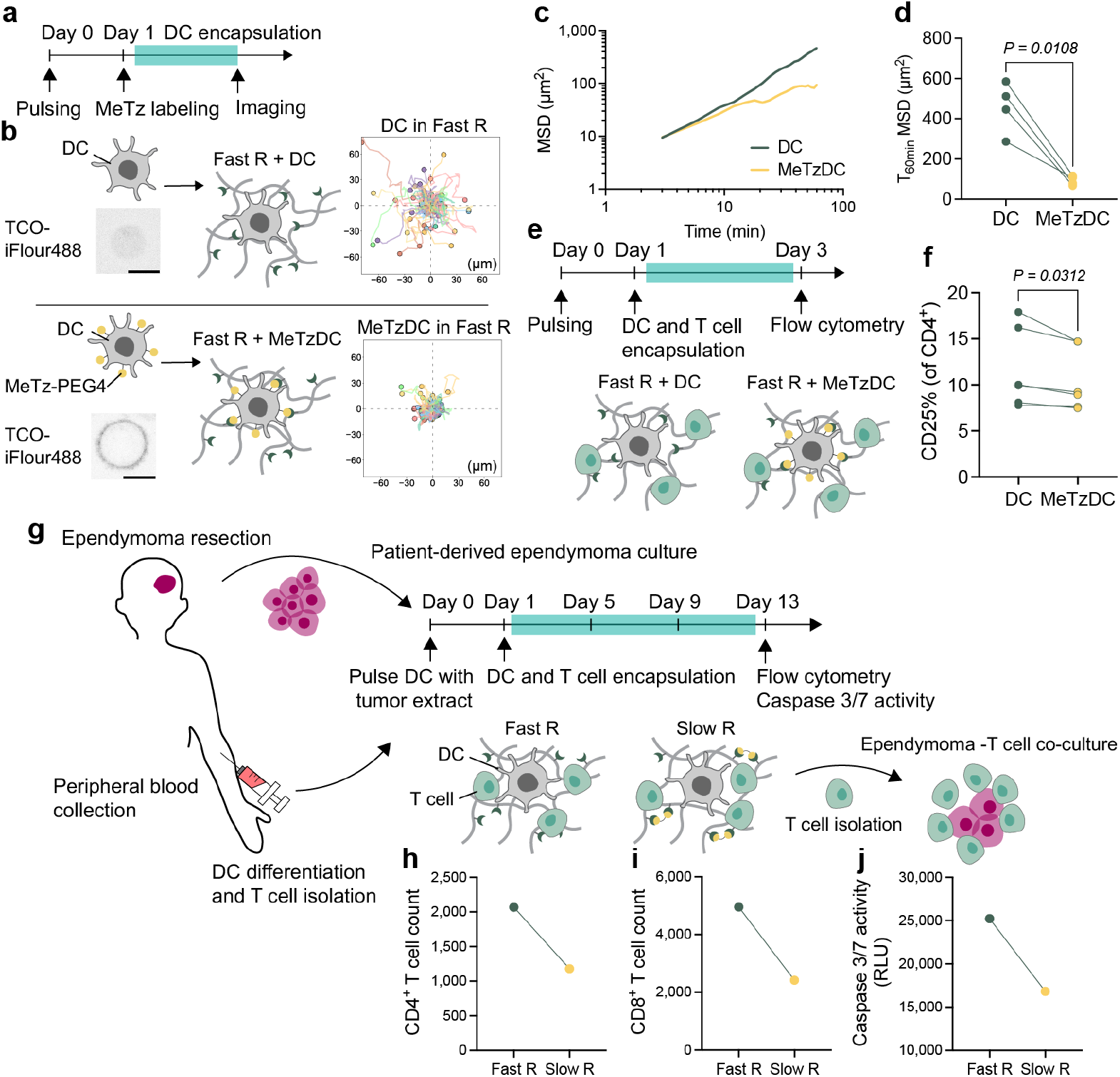
Confined DC migration in elastic ECM reduces T cell priming ability. **a**, Timeline for the conjugation of MeTz on DC surfaces. DCs were pulsed with SK-MEL5 lysates and activated with TNF-α. The following day, DCs were conjugated with MeTz via NHS chemistry and then encapsulated with T cells in Fast R gels for migration tracking. **b**, Schematic, representative image, and trajectory plot of unconjugated DCs or MeTzDCs, labeled by clicking with TCO-iFluor 488. Scale bar, 10 µm. **c**,**d**, Quantification of DC MSD over time, with MSD values at 60 min shown in paired donor samples; lines connect values from the same donor. *P* values were determined using a two-tailed paired t-test (*n* = 4). **e**, Timeline for co-encapsulation of DCs or MeTzDCs with T cells in Fast R gels. After pulsing with SK-MEL5 lysates and TNF-α activation, DCs or MeTzDCs were co-encapsulated with autologous T cells for 3 days, followed by flow cytometry analysis. **f**, Quantification of CD25 expression on CD4^+^ T cells co-cultured with DCs or MeTzDCs in Fast R gels; lines connect values from the same donor. *P* values were determined using a two-tailed Wilcoxon matched-pairs signed rank test (*n* = 6). **g**, Schematic and timeline of a patient autologous system with ependymoma cells, DCs, and T cells. Following ependymoma resection, PBMCs were isolated for DC differentiation and T cell isolation. DCs were pulsed with ependymoma lysates, activated with TNF-α, and encapsulated with T cells in 2 mg mL^−1^ Fast R or Slow R gels for 12 days. At day 12, gels were digested and T cells were co-cultured with ependymoma cells to assess their ability to induce cancer cell caspase-3/7 activity. **h**,**i**, Quantification of CD4^+^ and CD8^+^ T cell priming by DCs in Fast R or Slow R gels. **j**, Caspase-3/7 activity in ependymoma cells co-cultured with T cells primed by DCs in Fast R or Slow R gels. Data are from one patient; lines connect paired values from the same donor.

Finally, to assess the relevance of matrix viscoelasticity in a patient-derived setting, we established an autologous co-culture of ependymoma cells, DCs, and matched PBMCs from a pediatric patient (**Fig.6g**). After 12 days, both CD4^+^ and CD8^+^ T cells expanded more robustly in Fast R gels than in Slow R (**Fig.6h,i**), indicating impaired priming in the more elastic environment. Further, when T cells primed in Fast R or Slow R gels were subsequently co-cultured with ependymoma cells, cancer cells exposed to Slow R-primed T cells exhibited reduced caspase-3/7 activation, indicating weaker T cell cytotoxic activity (**Fig.6j**).

## Discussion

The findings of this study demonstrate that DC migration is markedly impaired in slow-relaxing (more elastic) extracellular matrices, due to diminished actomyosin-dependent collagen remodeling. Impaired migration in this setting emerged as a critical determinant in limiting DC-T cell interactions, reducing T cell priming, and weakening T cell responses. Prolonged confinement in slow-relaxing matrices induced a form of mechanomemory, with long-term deficits in migration, accompanied by altered chromatin accessibility at mechanosensing-related loci. Together, these findings identify viscoelasticity as a key regulator of DC function and point to mechanical confinement within tumors or fibrotic tissues as a barrier to effective immune activation.

DC migration was found to highly depend on actomyosin-driven collagen remodeling. In slow-relaxing matrices, where viscoelastic yielding is limited, contractile forces fail to generate persistent matrix remodeling, corresponding to impaired migration. This is consistent with prior work in DCs showing that actomyosin-generated contractility couples to central actin pools and WASp-driven cortical patches to deform restrictive matrices, thereby creating space for the nucleus and cell body to advance^14,47^. Importantly, inhibition of matrix metalloproteinases did not impair migration in fast-relaxing gels, whereas blocking actomyosin signaling abolished motility and remodeling, underscoring that path generation relies on cell-generated forces rather than proteolysis. Similar force-based strategies have been reported in monocytes and T cells^48,49^, suggesting that protrusive force generation is a conserved mechanism of immune cell migration that is particularly compromised in slow-relaxing ECM.

DC migration is essential for T cell priming, and our findings indicate that reduced motility in elastic ECM is the principal driver of impaired T cell proliferation and activation in this setting. This conclusion is consistent with intravital imaging studies showing that the frequency and stability of DC-T cell contacts within lymph nodes dictate the efficiency of activation^15,16,50^. Importantly, the diminished T cell activation by DCs in slow relaxing matrices effect cannot be explained by altered DC maturation, as expression of co-stimulatory and antigen-presenting molecules remained unchanged, nor by reduced cytokine output, which was in fact elevated in elastic matrices. Instead, our data argue that physical confinement of DCs limits their ability to remodel ECM and engage T cells, thereby uncoupling inflammatory activation from functional priming. These results underscore the central role of DC migration as a potential bottleneck for adaptive immune activation in more elastic environments.

Strikingly, DCs encapsulated in slow-relaxing matrices adopted a more inflammatory phenotype, consistent with reports that confinement-induced shape sensing triggers NF-κB signaling in immature DCs, leading to ARP2/3-dependent CCR7 upregulation and licensing of lymph node migration^51^. In our system, however, TNF-α-activated DCs did not further upregulate co-stimulatory molecules but instead amplified NF-κB/AP-1 signaling and inflammatory cytokine production. Similar effects have been described on stiff substrates, where LPS-stimulated DCs upregulate TLRs and secrete IL-6, as compared with soft substrates^52^. Parallel findings in other immune cells reinforce this view: monocytes in slow-relaxing elastic gels adopt a proinflammatory phenotype and preferentially differentiate into DCs^46^, while T cells show enhanced AP-1 activation via JNK signaling^41^. Although DCs in slow-relaxing gels secreted elevated IL-1β and IL-6, cytokines that support T cell survival^53,54^, this was insufficient to sustain T cell proliferation, and T cell activation was reduced, reflecting fewer productive DC-T cell encounters.

Prolonged encapsulation in slow-relaxing gels induced a form of mechanomemory, whereby DCs retained restricted migratory behavior even after transfer to fast-relaxing environments. This phenotype was associated with reduced chromatin accessibility at mechanosensing-related loci, consistent with recent studies showing that viscoelastic environments can durably remodel nuclear architecture and epigenetic state^40,41^. Such memory may compound the effects of confinement, as DCs not only migrate poorly within slow-relaxing matrices but may also carry forward impaired motility, limiting their ability to subsequently engage T cells or reach lymph nodes. In lymph nodes, productive priming relies on repeated, dynamic DC-T cell encounters^50,55^; by constraining migration and promoting long-lasting reprogramming, elastic regions of the TME may therefore act as local traps that diminish DC capacity to support immunity. Together, these findings establish ECM viscoelasticity as a key regulator of DC function and identify mechanical confinement as a barrier to effective antitumor immunity.

## Methods

### Collagen gel functionalization

The norbornene functionalization of collagen was performed as previously described^41^. In brief, Rat Tail Collagen Type I (Corning, 354236) was functionalized with 5-norbornene-2-acetic acid succinimidyl ester (nb-NHS) (Sigma Aldrich, 776173) at a ratio of 10 mg nb-NHS to 100 mg collagen. Collagen was first neutralized with NaOH to a pH of ∼7.5 and buffered with 10× phosphate-buffered saline (ThermoFisher, 70011044) to a concentration of 2 mg mL^−1^, followed by stirring at 4°C. Nb-NHS was dissolved in DMSO (Thermo Fisher, D12345) to an initial concentration of 10 mg mL^−1^ and then added to the neutralized collagen solution. The reaction proceeded for 4 hours at 4°C under rigorous stirring to delay collagen gelation. Afterward, 0.1 M acetic acid was added to quench the reaction and re-acidify the collagen solution. The product, norbornene-modified collagen, was dialyzed in 25 mM acetic acid for 3 days, filtered through a 0.45 μm filter (Corning, 430768), and lyophilized for future use.

### Rheological characterization

Rheological characterization of collagen gels was performed using a combined motor and transducer rheometer (AR-G2, TA Instruments) with a 25 mm cone-plate geometry. Neutralized collagen solution (100 µL) was loaded onto a 4 °C Peltier plate and allowed to gel at 37 °C for 30 min. After gelation, the gel was submerged in RPMI with DMSO or a MeTz-(PEG)5-MeTz crosslinker solution within a solution chamber to facilitate diffusion at 25 °C for 20 min. The solution was then removed, and the temperature was raised to 37 °C to initiate the click reaction, which proceeded for 30 minutes. An oscillatory frequency sweep (0.01 Hz to 5 Hz at 1% strain) was conducted at 37 °C to measure the storage modulus (G′), loss modulus (G″), and loss angle. A shear stress relaxation test was performed by applying 10% shear strain and maintaining the strain while recording changes in shear stress over 2 hours at 37 °C. A humidity chamber was used throughout the experiments to prevent dehydration.

### DC differentiation and T cell isolation

De-identified apheresis collars were obtained from Brigham and Women’s Hospital. Human PBMCs were then isolated using a Ficoll gradient (Cytiva, 17144002). Human monocytes were isolated from PBMCs 6 days prior to the experiments using a magnetic bead-based CD14^+^ isolation kit (Miltenyi, 130-050-201). Following isolation, the monocytes were differentiated into DCs by culturing them in DC differentiation media, consisting of RPMI 1640 medium (ThermoFisher, 11875093) supplemented with 10% heat-inactivated fetal bovine serum (ThermoFisher, 10082147), 1% penicillin-streptomycin (ThermoFisher, 15140122), 50 ng mL^−1^ IL-4 (PeproTech, 200-04), and 50 ng mL^−1^ GM-CSF (PeproTech, 300-03). After 5 days of culturing, 100 ng mL^−1^ TNF-α (PeproTech, 300-01A) was added to the media and incubated for 24 hours to activate the DCs. Flow cytometry confirmed DC differentiation and upregulation of activation markers in response to TNF-α (**Supplementary Fig. 12**). Pan T cells were isolated from PBMCs using a magnetic bead-based Pan T cell isolation kit (Miltenyi, 130-096-535) following the manufacturer’s protocol.

### Cell encapsulation and collagen crosslinking

Lyophilized nb-Col was first dissolved at 6 mg mL^−1^ in 25 mM acetic acid at 4 °C for 3 days. To cast fast-relaxing (non-click crosslinked) and slow-relaxing (click crosslinked) gels, 6 mg mL^−1^ nb-Col was mixed on ice with 10x PBS, Milli-Q water, and neutralized with 1M NaOH to achieve a final pH of ∼7.5 and a final collagen concentration of 2.2 mg mL^−1^ or 4.4 mg mL^−1^. Activated DCs were resuspended in PBS (ThermoFisher, 14190235) on ice, and 100,000 cells were mixed with the collagen solution at a 1:10 volume ratio to reach a final collagen concentration of 2 mg mL^−1^. The collagen-DC solution was then added to the 10-mm-diameter center well of a 12-well MatTek plate (P12G-1.5-10-F) or a 24-well MatTek plate (P24G-1.5-10-F) and incubated at 37 °C for 30 min to allow collagen gelation. After gelation, the plate was transferred to room temperature for 10 minutes. Methyltetrazine-(PEG)5-methyltetrazine was dissolved in 1 mL RPMI media to a final concentration of 282 µM and added to the gel for crosslinking. DMSO, at the same volume as the crosslinker, was diluted in RPMI and added as a control. After 10 minutes of incubation at room temperature to allow the crosslinker to diffuse within the collagen while minimizing crosslinking, the solution was removed, and the plate was transferred to 37 °C for further crosslinking and incubated for another 30 minutes. Following crosslinking, collagen gels underwent three 10-minute washes with RPMI media to remove excess crosslinker and were cultured in DC differentiation media supplemented with 100 ng mL^−1^ TNF-α.

### DC and T isolation from collagen gels

To isolate DCs from collagen gels, the gels were transferred to an RPMI solution supplemented with 10% FBS and 80 U mL^−1^ collagenase type IV (Thermo Fisher, 17104019) and incubated at 37 °C for at least 30 minutes. During the incubation, the collagen gel was mechanically disrupted in collagenase solution by pipetting through 1 mL pipette tips every 10 minutes until the collagen gel was fully digested. Afterward, an equal volume of EasySep buffer (Stemcell Technologies, 20144) was added to the solution to stop the reaction. The cell solution was then collected, washed twice with PBS, and resuspended for downstream analysis.

### Cancer cell culture and lysate preparation

SK-MEL-5 cells were cultured in DMEM (Gibco, 11965092) supplemented with 10% heat-inactivated fetal bovine serum and 1% penicillin–streptomycin. The ependymoma cell line was maintained in human NeuroCult NS-A proliferation medium (StemCell Technologies, 05751) supplemented with 1:100 Antibiotic–Antimycotic (Gibco, Cat#15240112), 1:100 HEPES (Life Technologies, Cat#15630080), 20 ng mL^−1^ hEGF (Shenandoah Biotechnology, Cat#100-26AF), 20 ng mL^−1^ hFGF (Shenandoah Biotechnology, Cat#100-146AF), and 1:1000 heparin (StemCell Technologies, 07980), and cultured in ultra-low-adherent 6-well plates (Corning, 3471). Both cell lines were maintained in a 37 °C incubator with 5% CO_2_. For tumor lysate preparation (SK-MEL-5 and ependymoma), cells were washed twice with PBS and incubated with TrypLE Express (for SK-MEL-5; Gibco, 12604013) or Accutase (for ependymoma; StemCell Technologies, 07920) for 5 min at 37 °C to detach adherent cells or dissociate clumps. The enzymatic reaction was quenched by adding complete medium, and cells were pelleted by centrifugation at 350 × g for 5 min. Tumor lysates were generated by four freeze–thaw cycles, alternating between dry ice and a 37 °C water bath. Cell debris was removed by centrifugation at 14,000 × g for 10 min at 4 °C, and protein concentration was determined using the Pierce BCA Protein Assay Kit (Thermo Scientific, A65453) according to the manufacturer’s instructions. Lysates were aliquoted and stored at –80 °C until use.

### DCs-T cell co-culturing

After differentiation, DCs were pulsed with 200 μg mL^−1^ SK-MEL-5 tumor extract, prepared by four freeze-thaw cycles, for 3 hours in RPMI 1640 supplemented with 1% penicillin–streptomycin, followed by activation with TNF-α for 16 hours. Subsequently, DCs and T cells labeled with CFSE or CellTrace Deep Red were co-encapsulated at a 1:10 (DC:T) ratio in either fast- or slow-relaxing collagen gels, as previously described. Encapsulated cells were cultured in X-VIVO 15 medium (Lonza, 02-053Q) supplemented with 10 ng mL^−1^ IL-2 (PeproTech, 200-02), 10 ng mL^−1^ IL-7 (PeproTech, 200-07), and 10 ng mL^−1^ IL-15 (PeproTech, 200-15). Media were refreshed every 4 days or earlier as needed. After 12 days, collagen gels were digested as described previously, and samples were collected for downstream analysis.

### Transwell assay

After pulsing DCs with SK-MEL-5 tumor lysates and activating them with TNF-α, DCs and T cells were co-encapsulated in either Fast R or Slow R gels in the upper chambers of Transwell inserts with polycarbonate membranes (3 μm pore size; Corning, 3415), which allow T cell migration. On day 5, the inserts were transferred to fresh lower chambers containing 25,000 SK-MEL-5 tumor cells per well. Cultures were maintained for an additional 7 days. On day 12, collagen gels from the upper chambers were digested as described previously. Cells in the lower chambers were harvested directly without digestion for subsequent analysis.

### Patient samples

Patient-derived cell line and peripheral blood samples were obtained from ependymoma patients undergoing surgical resection at Boston Children’s Hospital, after receiving informed consent and approval from the institutional review board at Dana Farber Cancer Institute (DFCI 10-417). Patient-derived cell line was used for preparation of tumor lysates or for co-culture assays as indicated. PBMCs were isolated from whole blood by density-gradient centrifugation (Ficoll-Paque PLUS, Cytiva) and used for autologous DC differentiation and T cell isolation. All procedures were performed in accordance with ethical guidelines and approved protocols.

### Time-lapse Imaging

After encapsulation, DCs were incubated in DC differentiation media supplemented with 100 ng mL^−1^ TNF-α at 37 °C for 1 hour prior to imaging. For tracking DC migration, SiR-DNA (Cytoskeleton, CY-SC007) was added 1 day prior to imaging at a final concentration of 200 nM to label cell nuclei. For small molecule inhibition, the following inhibitors were individually added to the DC differentiation media 1 hour before and during time-lapse imaging: 25 µM para-amino-blebbistatin (Cayman, 22699), 10 µM Y27362 (Stem Cell Technologies, 72302), 10 µM marimastat (ThermoFisher, J67288.MCR), 10 µM GM6001 (Sigma-Aldrich, 364206), 1 μM IPI-549 (Selleckchem, S8330), 10 µM defactinib (Abcam, ab254452), and 500 nM latrunculin A (Abcam, ab144290). DMSO at an equivalent volume was included as a control. Time-lapse imaging was performed using Zeiss LSM 980 and Leica Stellaris confocal microscopes, with images captured every 3 minutes over a 2-hour span. Images were processed using FIJI software^56^, and tracking was conducted with TrackMate^57^. Cell migration trajectory plots and mean squared displacement analyses were performed using custom Python scripts. For tracking collagen displacement, collagen was fluorescently labeled by incubating DC-encapsulating Nb-Col with tetrazine-iFluor 488 (AAT Bioquest, 1014) at 37 °C for 1 hour to allow the click reaction, followed by two washes with DC media. SiR-DNA (Cytoskeleton, CY-SC007) was added 1 day prior to imaging at a final concentration of 200 nM to label cell nuclei, and SPY555-FastAct (Cytoskeleton, CY-SC205; 1:2000) was added 1 hour prior to imaging to label actin. Time-lapse imaging for collagen displacement was performed using a Zeiss LSM 980 confocal microscope, with images captured every minute over a 10-minute span. Particle image velocimetry was performed using the MATLAB-based PIVlab toolbox^58^. Collagen fibrils labeled with tetrazine-iFluor 488 were imaged by time-lapse confocal microscopy, and displacement fields were calculated between consecutive frames. Vector fields were validated and smoothed using standard PIVlab filters, and displacement magnitudes were summed over the 10-min imaging period to quantify total collagen remodeling. For visualization, displacement maps were color-coded according to magnitude, and the primary axis of collagen displacement was determined for downstream analysis.

### Flow cytometry staining

Prior to flow cytometry staining, DCs and T cells were isolated from collagen gels by collagen digestion as previously described, followed by rinsing the wells with PBS and collecting the rinse. The cells were washed with PBS, then stained for dead cells using the dead cell staining kit (Invitrogen, L34982 or L23105) for 10 minutes at 4°C. After staining, the cells were washed twice with FACS buffer (Invitrogen, 00-4222-26). For blocking, human Fc block (BioLegend, 422302) and Brilliant Stain Buffer (BD, 563794) were added to the cells and incubated at 4°C for 10 minutes. Subsequently, the corresponding antibodies, diluted 1:100, were directly added to the cell solution. After a 30-minute incubation at 4°C, the cells were washed twice with FACS buffer, resuspended in FACS buffer, and analyzed using a Sony ID7000. Gating was performed using fluorescence-minus-one controls. The gating strategy for CD4^+^ and CD8^+^ T cells and DCs is shown in **Supplementary Fig.13**. The complete list of antibodies used for flow cytometry is provided in **Supplementary Table 1**.

### Cytokine secretion

TNF-α-activated DCs were encapsulated in either fast- or slow-relaxing collagen gels in 12-well plates and cultured for 3 days in DC differentiation medium supplemented with 100 ng mL^−1^ TNF-α. On day 3, culture supernatants were collected and centrifuged at 3,000 x g for 10 min to remove debris, then stored at −80 °C for subsequent ELISA analysis. For ELISA, plates were prepared according to the manufacturer’s instructions (R&D Systems, DY201 and DY206), and signal was detected using a microplate reader (BioTek Synergy H1, Agilent).

### Pore size imaging and measurement

Fast R or slow R matrices were fluorescently labeled by incubation with 10 μM tetrazine-iFluor 488 in PBS for 1 h at 37 °C, followed by three PBS washes. Imaging was performed on a Zeiss LSM 980 confocal microscope. Z-stacks were acquired at 0.24 μm intervals over a total thickness of approximately 5.3 μm (22 slices), and maximum intensity projections were generated to obtain a 2D representation of the collagen fibril network. Pore size was quantified from the projected images using Fiji. Images were thresholded, binarized, and inverted to segment the pores. The area of each pore was measured using the Analyze Particles function, and the equivalent diameter was calculated as 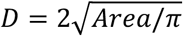, representing the diameter of a circle with the same area as the pore.

### MeTz conjugation on DCs

After pulsing and activation, DCs were washed twice with PBS and incubated with 25 µM methyltetrazine-PEG4-NHS ester (Broadpharm, Cat#BP-22945) for 30 min at 4 °C. The reaction was quenched by adding FBS to a final concentration of 20%. After two washes, DCs were encapsulated in Fast R or unmodified collagen, with or without T cells, for flow cytometry analysis or migration assays, respectively. To monitor MeTz retention on DCs, MeTz-labeled cells were maintained in DC differentiation medium supplemented with TNF-α and collected at the indicated time points. Cells were stained with 10 µM TCO-iFluor 488 (AAT Bioquest, 1005) for 30 min at 37 °C, followed by two PBS washes. Images were acquired on a Leica Stellaris confocal microscope.

### Caspase-3/7 cytotoxicity assay

The cytotoxic activity of T cells was quantified using the Caspase-Glo® 3/7 assay (Promega, G8091) in 96-well white opaque plates. T cells derived from Fast R or Slow R conditions were collected and added to cancer cells at an effector-to-target ratio of 3:1. Co-cultures were established by adding 50 µL of the T-cell suspension to wells containing cancer cells. Cultures were incubated for 4 h at 37 °C, equilibrated to room temperature, and then treated with 100 µL of caspase-3/7 reagent. After a 30-min incubation at room temperature, luminescence was measured using a microplate reader (Synergy H1, BioTek; Agilent) in luminescence mode.

### Sample preparation for bulk RNA sequencing and ATAC sequencing

Monocytes isolated from four or three donors were differentiated to DC and activated with TNF-α, as described, for bulk RNA sequencing and ATAC sequencing, respectively. The resulting DCs were then encapsulated in fast- and slow-relaxing 2 mg mL^−1^ collagen gels for 3 days. Afterward, DCs were isolated from the collagen gels by collagenase digestion, as described. From each sample, 50,000 cells were used for ATAC-seq library preparation using the Zymo-Seq ATAC Library Kit (Zymo Research, D5458). The remaining cells were used for RNA isolation with the PureLink RNA Micro Scale Kit (Thermo Fisher, 12183016), following the manufacturer’s instructions.

### Analysis for bulk RNA sequencing and ATAC sequencing

For principal component analysis, raw gene counts were log-transformed using log1p to reduce the influence of highly expressed transcripts, transposed, and standardized across samples; PCA was performed with scikit-learn (Python). Differential expression analysis was carried out with DESeq2 using variance-stabilizing transformation for normalization, and genes with an adjusted *P* < 0.05 and |log_2_ fold change| > 1 were considered significant. Gene Ontology enrichment was performed with Enrichr (gseapy), and selected terms were visualized. Heatmaps were generated from z-score–normalized expression data. Transcription factor activity was inferred using DoRothEA, and volcano plots were generated to display log_2_ fold change against –log_10_ adjusted *P*. For ATAC-seq, reads were adapter-trimmed with Cutadapt (v5.0), aligned to hg19 with BWA-MEM2 (v2.2.1), sorted with SAMtools (v1.21), and deduplicated with Picard (v3.3.0). Peaks were called with MACS2 (v2.2.9.1) using parameters –nomodel –shift −100 –extsize 200 –qvalue 0.01. Differential accessibility was assessed with DiffBind using a linear model (∼ Replicate + Condition). Peaks were annotated with ChIPseeker (TxDb.Hsapiens.UCSC.hg19.knownGene) to assign genomic context and nearest genes within ±3 kb of the transcription start site. Chromatin accessibility around transcription start sites was quantified and visualized with deepTools (v3.5.6).

### Statistical analysis

Statistical testing was performed using GraphPad Prism (v.10), with detailed methods described in the text and figure legends. Data were first assessed for normality to determine whether parametric or nonparametric analyses were applied. *P* < 0.05 was considered significant.

## Acknowledgments

We thank T. Harimoto and K. Adu-Berchie for their scientific input, and T. Ferrante for support with microscope imaging. W.-H.J. was supported by the National Cancer Institute (K00CA253759). Y.B. was supported by the National Cancer Institute (T32CA136432). Bulk RNA-seq and ATAC-seq were performed at the Bauer Core Facility at Harvard University. Additional support was provided by the Wellcome Leap HOPE program.

## Contributions

W.-H.J. and D.J.M. conceptualized and designed the study. W.-H.J., E.H., A.H. and S.I. performed experiments and analyzed data. J.M.P. analyzed sequencing data. Y.B. provided guidance on study design and contributed patient samples. W.-H.J. and D.J.M. wrote the manuscript. All authors reviewed and edited the manuscript.

## Competing interests

D.J.M. has sponsored research, consults and/or has stock options/stock in Medicenna, Lyell, Attivare, Epoulosis, Limax Biosciences, Lightning Bio and Oddity Tech; has licensed intellectual property with Alkem and Amend Surgical; and is on the board of directors for ATCC. The other authors declare no competing interests.

## Data availability

The data supporting the findings of this study are included in the paper and its Supplementary Information. Bulk RNA-seq and ATAC-seq data have been deposited in the NCBI Gene Expression Omnibus under accession codes GSE309231 and GSE309232. Additional data are available from the corresponding author upon reasonable request.

