## Supplementary Information for "Matrix viscoelasticity regulates dendritic cell migration and immune priming"

This PDF file includes:

Supplementary Figs.1-13

Legends for Supplementary Movies 1-4

Supplementary Table 1

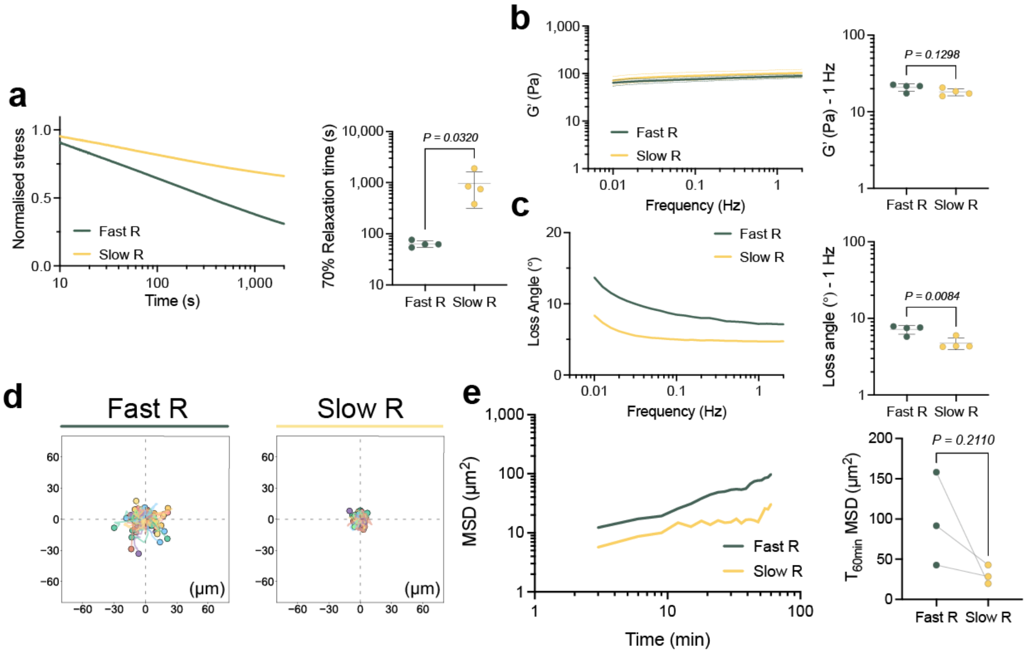


**Supplementary Fig.1 | Human DC migration is confined in the 4 mg mL$^{-1}$ slow-relaxing collagen gels.**

**a**, Stress relaxation profiles of 4 mg mL$^{-1}$ Fast R and Slow R gels, and quantification of the time required for stress to decay to 70% of the initial value. **b**, Storage modulus (G') of 4 mg mL$^{-1}$ Fast R and Slow R gels determined by frequency sweep, and corresponding quantification at 1 Hz. **c**, Loss angle of 4 mg mL$^{-1}$ Fast R and Slow R gels determined by frequency sweep, with quantification at 1 Hz. Results in **a–c** are from four independently formed gels and are presented as mean ± s.d. *P* values were determined using a two-tailed unpaired t-test. **d,** Representative migration trajectories of DCs encapsulated in 4 mg mL$^{-1}$ Fast R or Slow R gels over 60 minutes. **e**, Quantification of DC MSD over time within Fast R or Slow R gels, with MSD values highlighted at the 60-minute time point. Data are representative of three independent donors; lines connect values from the same donor. *P* values were determined using a two-tailed paired t-test ($n$ = 3).

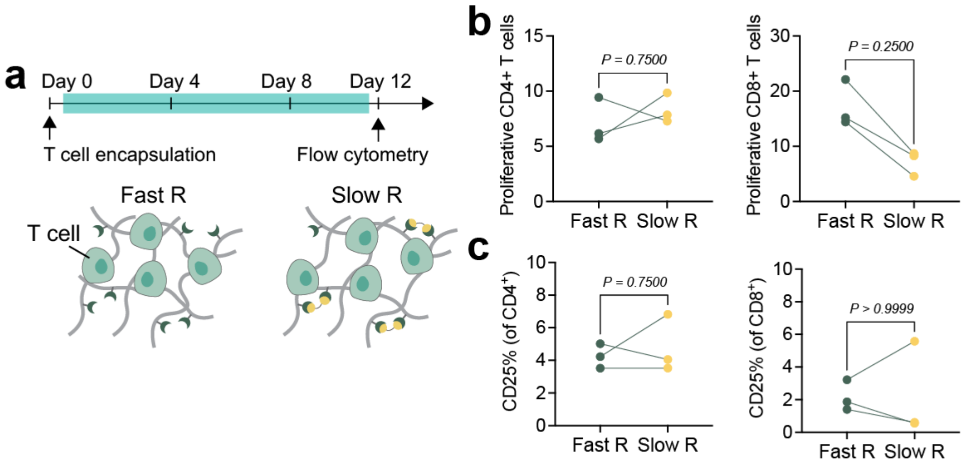


**Supplementary Fig.2 | T cell proliferation and activation in monoculture are comparable in Fast R and Slow R matrices.**

**a**, Schematic and timeline of unactivated T cells encapsulated in 2 mg $mL^{-1}$ Fast R or Slow R matrices for 12 days. **b**, Quantification of proliferating $CD4^{+}$ and $CD8^{+}$ T cells, measured as reduction in deep red dye signal. **c,** Quantification of CD25+ $CD4^{+}$ and $CD8^{+}$ T cells, as measures of activation. Data are representative of results from three independent donors; lines connecting data points in the Fast R and Slow R groups indicate cells derived from the same donor. *P* values were determined using a two-tailed Wilcoxon matched-pairs signed rank test ($n$ = 3).

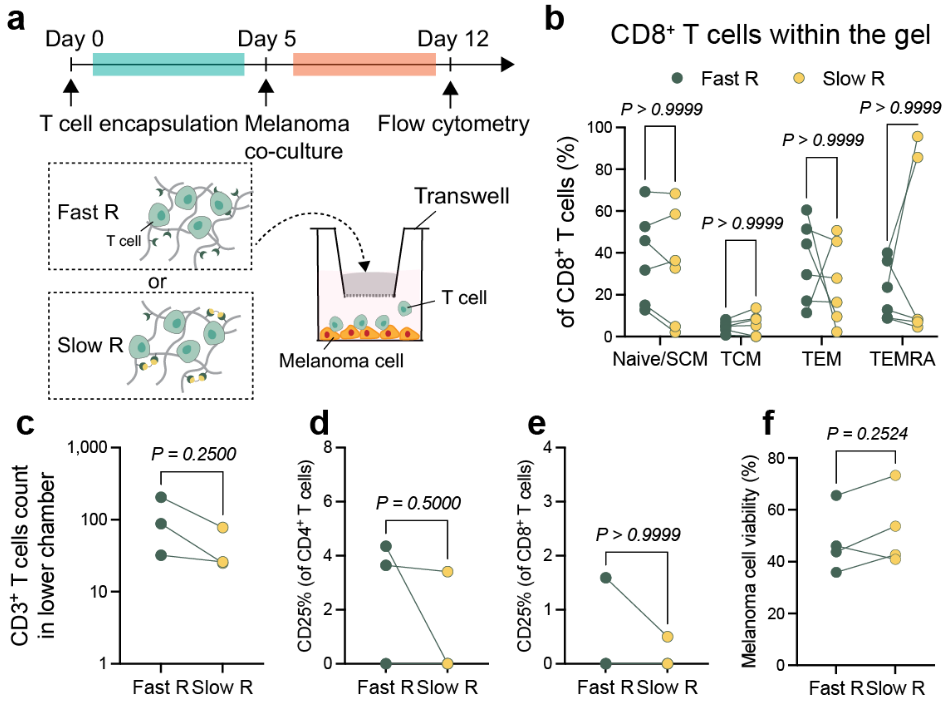


**Supplementary Fig.3 | T cell subset differentiation, activation, and migration are comparable in Fast R and Slow R matrices in the transwell assay, in absence of DCs.** **a**, Schematic and timeline of unactivated T cells encapsulated in 2 mg $mL^{-1}$ Fast R or Slow R matrices for 12 days in a transwell assay; at day 5, inserts were transferred to wells containing melanoma cells (SK-MEL5). **b**, Quantification of $CD8^+$ T cell subsets within the gel after 12 days of culture, defined by CD45RA and CCR7 expression (Naïve/SCM: $CCR7^+CD45RA^+$, TCM: $CCR7^+CD45RA^-$, TEM: $CCR7^-CD45RA^-$, TEMRA: $CCR7^-CD45RA^+$). *P* values were determined using a two-way repeated-measures ANOVA ($n$ = 6 donors, each measured across all conditions), followed by Šídák's multiple comparisons test to assess differences between Fast R and Slow R for each T cell subset. **c**, Quantification of $CD3^+$ T cells that migrated to the bottom chamber after 7 days of co-culture with melanoma cells. **d,e**, Quantification of $CD4^+$ and $CD8^+$ T cells expressing CD25 following migration through the transwell. **f**, Viability of melanoma cells after 7 days of co-culture with T cells. Data are representative of six donors (**b**) or three to four donors (**c–f**); lines connect paired data points from the same donor in the Fast R and Slow R groups. *P* values for (**c–e**) were determined using a two-tailed Wilcoxon matched-pairs signed-rank test, and the *P* value for (**f**) was determined using a two-tailed paired t-test ($n$ = 3 for **c–e**; $n$ = 4 for **f**).

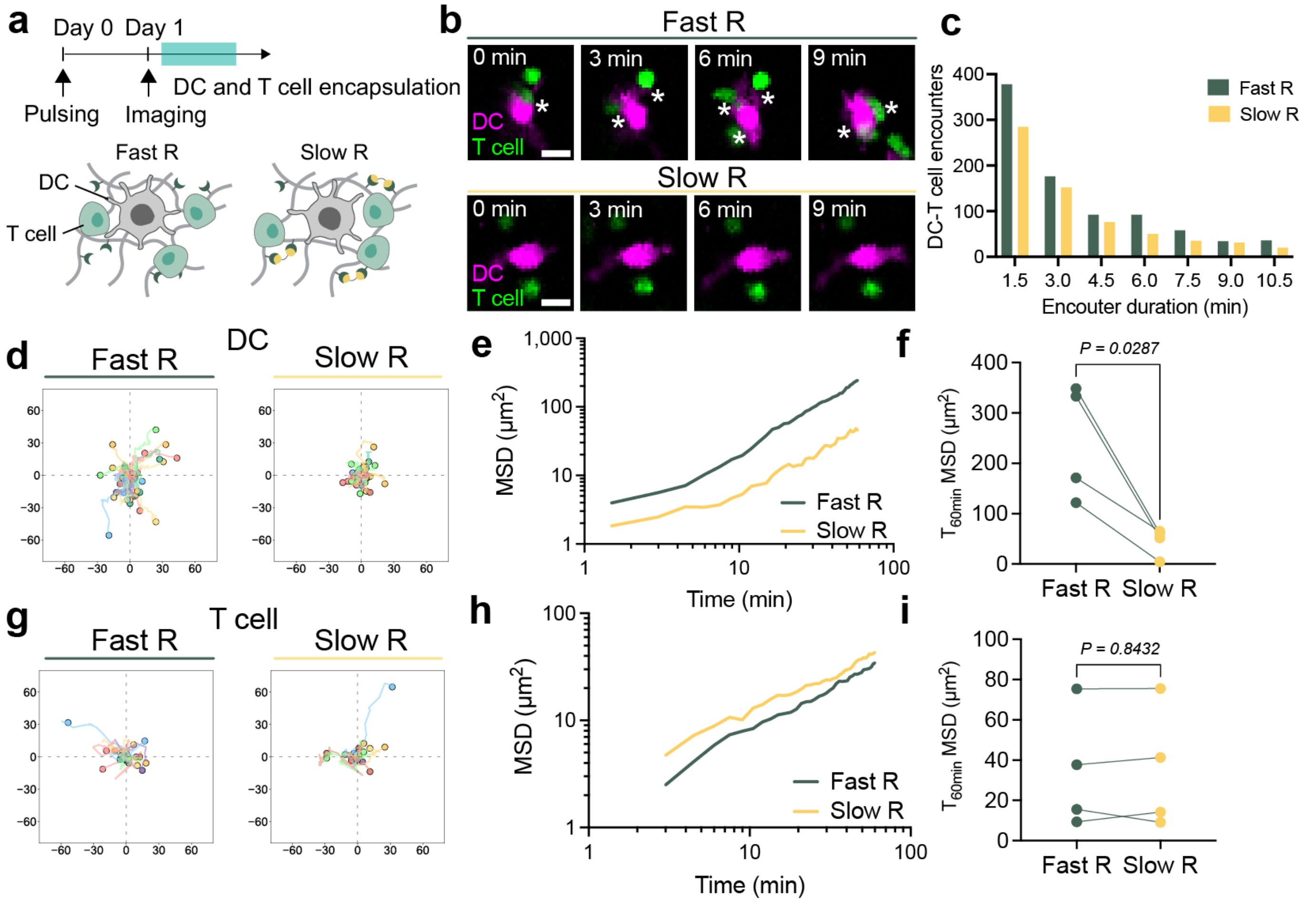


**Supplementary Fig.4 | Confined DC migration reduces DC–T cell interactions.**

**a**, Schematic and timeline of co-culturing DCs and unactivated T cells in 2 mg $mL^{-1}$ Fast R or Slow R matrices. DCs were pulsed with SK-MEL5 tumor extract and activated with TNF-α for 16 hr before encapsulation with autologous T cells. **b**, Representative images of DCs (magenta, CellTrace Yellow) and T cells (green, Deep Red Cell Tracer) in Fast R or Slow R matrices. Asterisks denote DC–T cell encounters. **c**, Histogram of DC-T encounters, as a function of duration, pooled from five independent donors. **d**, Representative trajectories of DCs co-encapsulated with T cells in 2 mg $mL^{-1}$ Fast R or Slow R gels over 60 min. **e,f**, Quantification of DC MSD over time, with MSD values at 60 min shown. **g**, Representative trajectories of T cells co-encapsulated with DCs in 2 mg $mL^{-1}$ Fast R or Slow R gels over 60 min. **h,i**, Quantification of T cell MSD over time, with MSD values at 60 min shown. *P* values were determined using a two-tailed paired t-test ($n$ = 4); lines connect paired data points from the same donor in the Fast R and Slow R groups.

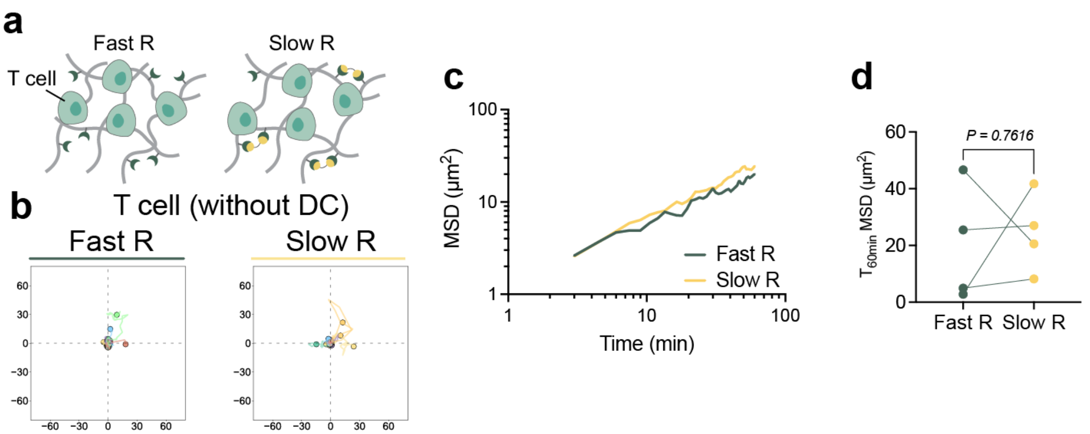


**Supplementary Fig.5 | Migration of unactivated T cells is not affected by ECM viscoelasticity.**

**a**, Schematic and timeline of unactivated T cells encapsulated in 2 mg $mL^{-1}$ Fast R or Slow R matrices. **b**, Representative trajectories of T cells cultured without DCs in Fast R or Slow R gels over 60 min. **c,d**, Quantification of T cell MSD over time, with MSD values at 60 min shown. Data are from four independent donors; lines connect paired data points from the same donor in the Fast R and Slow R groups. *P* values were determined using a two-tailed paired t-test ($n$ = 4).

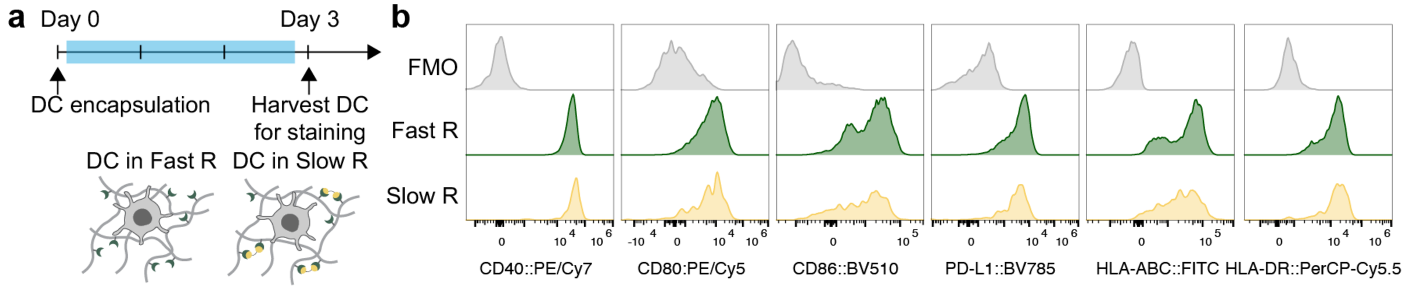


**Supplementary Fig.6 | Expression of costimulatory and antigen-presenting molecules on DCs remains comparable in fast- or slow-relaxing matrices.**

**a,** Schematic and timeline of TNF-α–activated DCs encapsulated in 2 mg $mL^{-1}$ Fast R or Slow R matrices. **b**, Representative histograms of surface marker expression after 3 days of encapsulation, selected from four independent donors. Fluorescence minus one (FMO) controls include all antibodies except the corresponding marker antibody.

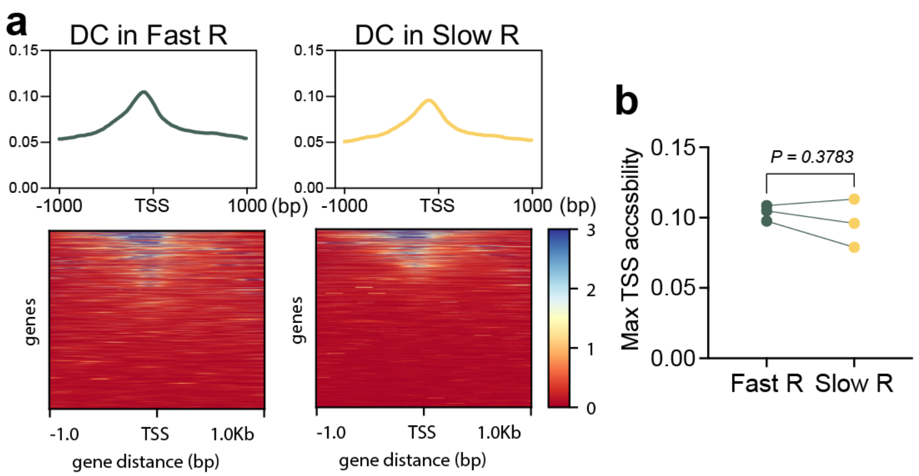


**Supplementary Fig.7 | Chromatin accessibility at transcription start sites (TSS) remains comparable in DCs encapsulated in Fast R or Slow R matrices.**

**a**, ATAC-seq signal profiles (top) showing normalized read density (fragments per million mapped reads, FPM) across ±1 kb from the TSS of the top 1,000 genes, and heatmaps (bottom) showing normalized accessibility ($log_2$ read density) for DCs cultured in 2 mg $mL^{-1}$ Fast R or Slow R matrices. **b**, Quantification of maximum TSS accessibility (normalized ATAC-seq read density, FPM) in paired donor samples; lines connect values from the same donor. *P* values were determined using a two-tailed paired t-test (*n* = 3).

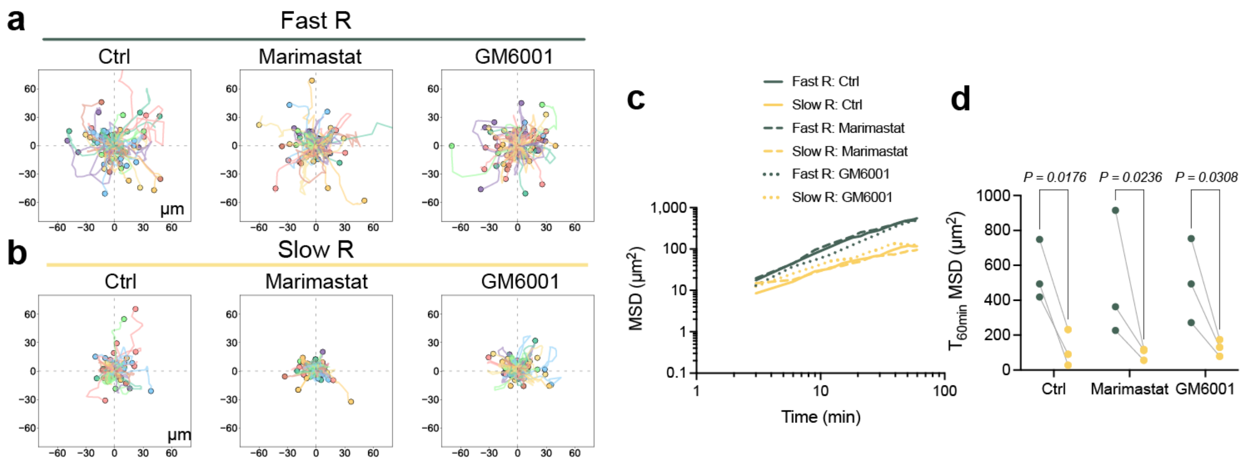


**Supplementary Fig.8 | DC migration is independent of MMP activity.**

**a,b,** Representative trajectories over 60 min of DCs treated with DMSO (Ctrl), Marimastat, and GM6001, with encapsulation in 2 mg $mL^{-1}$ Fast R or Slow R gels. **c,d**, Quantification of DC MSD over time, with MSD values at 60 min shown in paired donor samples; lines connect values from the same donor. *P* values were determined using a two-way repeated-measures ANOVA with uncorrected Fisher's LSD, with viscoelasticity and treatment as factors, and each donor measured across all conditions ($n$ = 3).

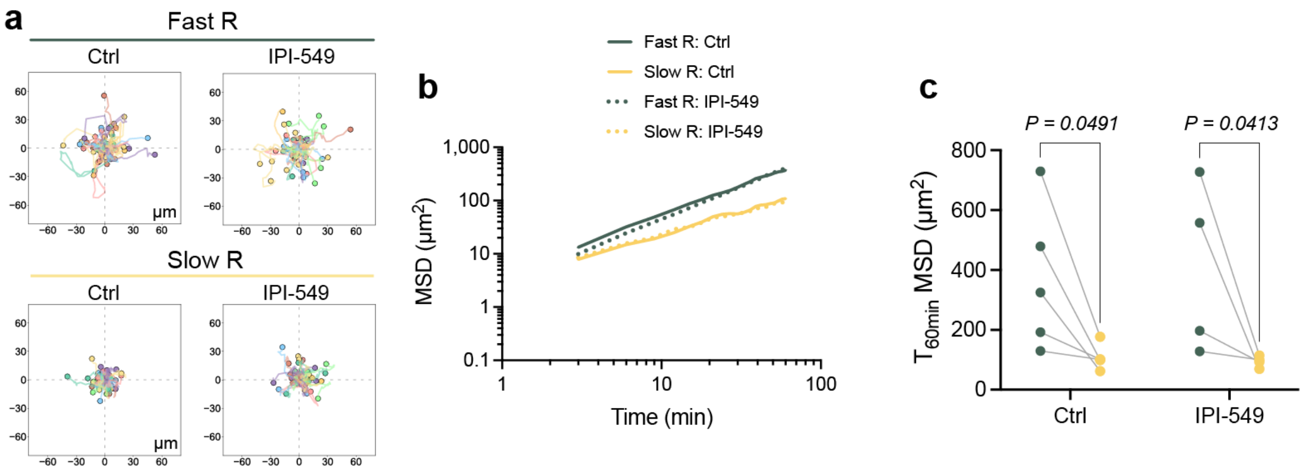


**Supplementary Fig.9 | DC migration is independent of PI3K activity.**

**a,b,** Representative trajectories over 60 min of DCs treated with DMSO (Ctrl) or IPI-549 and encapsulated in 2 mg mL$^{-1}$ Fast R or Slow R gels. **c,d**, Quantification of DC MSD over time, with MSD values at 60 min shown in paired donor samples; lines connect values from the same donor. *P* values were determined using a two-way repeated-measures ANOVA with uncorrected Fisher's LSD, with viscoelasticity and treatment as factors, and each donor measured across all conditions ($n$ = 5 for Ctrl, $n$ = 4 for IPI-549).

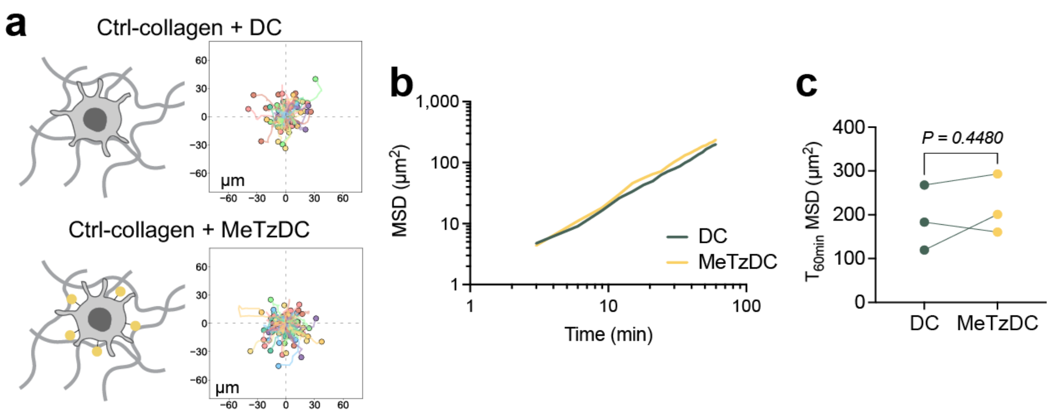


**Supplementary Fig.10 | MeTz conjugation does not alter DC migration.**

**a,** Schematic and representative trajectories of untreated DCs or MeTz-conjugated DCs (MeTzDCs) encapsulated in unmodified collagen (Ctrl-collagen) over 60 min. **b,c,** Quantification of DC MSD over time, with MSD values at 60 min shown in paired donor samples; lines connect values from the same donor. *P* value was determined using a two-tailed paired t-test ($n$ = 3).

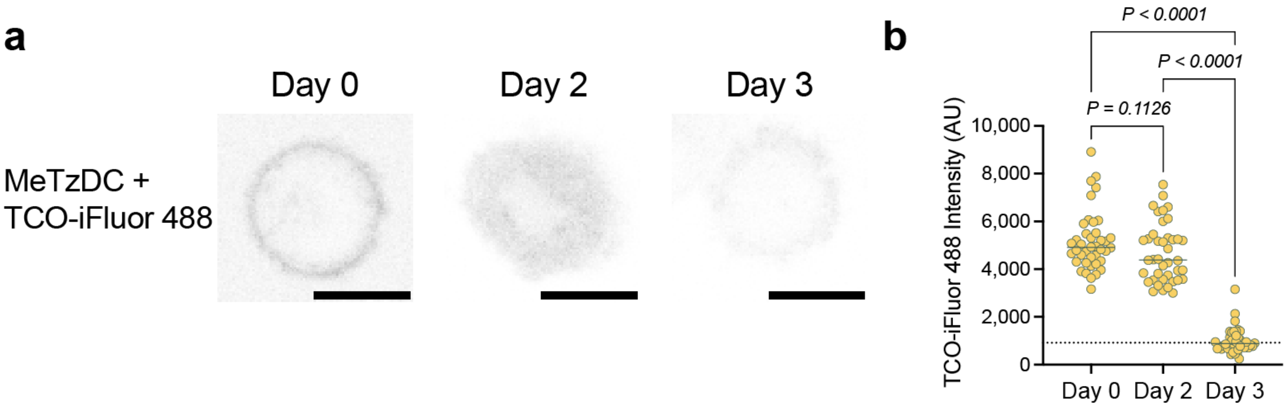


**Supplementary Fig.11 | Duration of MeTz retention on DCs.**

**a**, Representative images of MeTz-conjugated DCs clicked with TCO-iFluor 488 at day 0, day 2, and day 3. Scale bar, 10 μm. **b**, Fluorescence intensity of DCs after TCO-iFluor 488 staining at day 0, day 2, and day 3. The dotted line denotes the background signal from unconjugated DCs stained with TCO-iFluor 488. Lines indicate the median. *P* values were determined using an ordinary one-way ANOVA with Tukey's multiple-comparisons test. Each dot represents a single cell (40 cells total for each time point) from one donor.

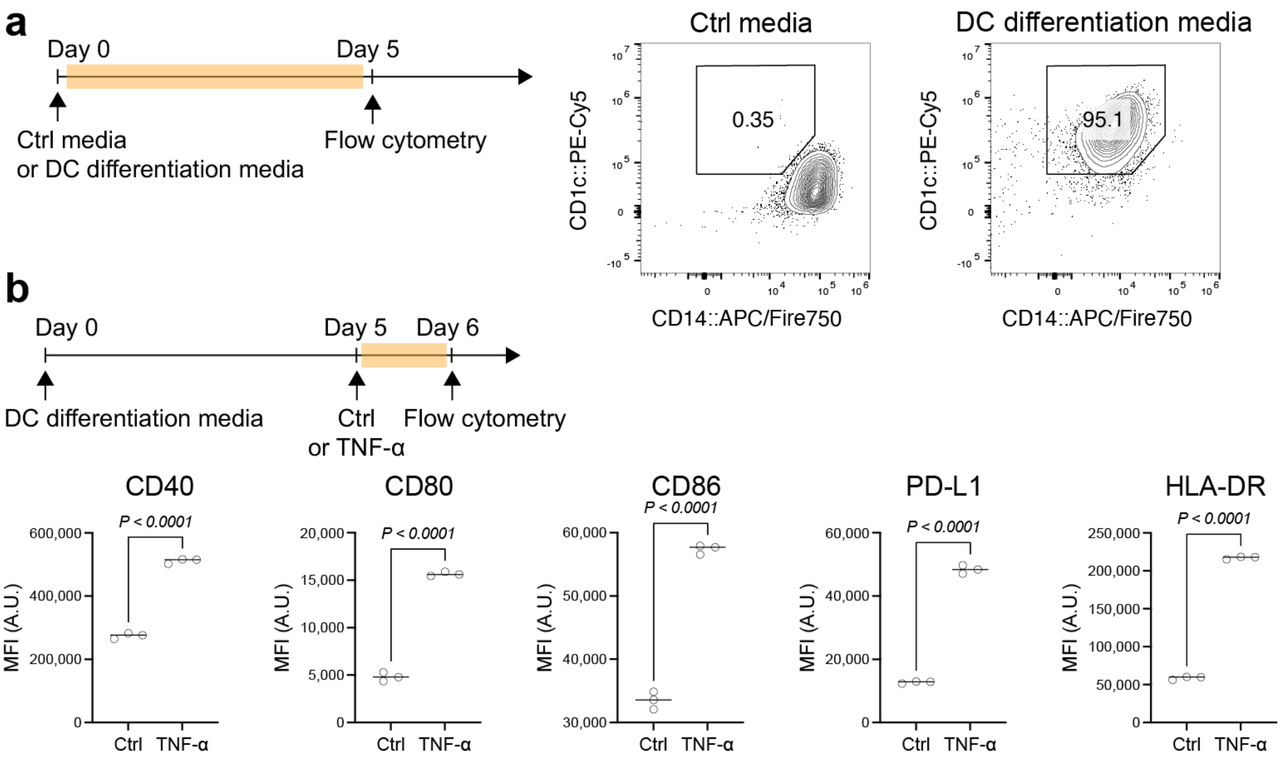


**Supplementary Fig.12 | Validation of DC differentiation and activation.**

**a**, Timeline and representative flow cytometry plots demonstrating human DC differentiation. Monocytes were cultured in either control medium or DC differentiation medium; the control medium contained all components except IL-4 and GM-CSF. After 5 days, cells were harvested for flow cytometry. DC differentiation was defined as loss of CD14 and gain of CD1c ($CD14^{-}CD1c^{+}$). **b**, Timeline and mean fluorescence intensity quantification of DCs cultured in differentiation medium without (Ctrl) or with TNF-α to assess activation (CD40, CD80, CD86, PD-L1, HLA-DR). *P* values were determined using an unpaired t-test. Each dot represents one of three technical replicates from a single donor.

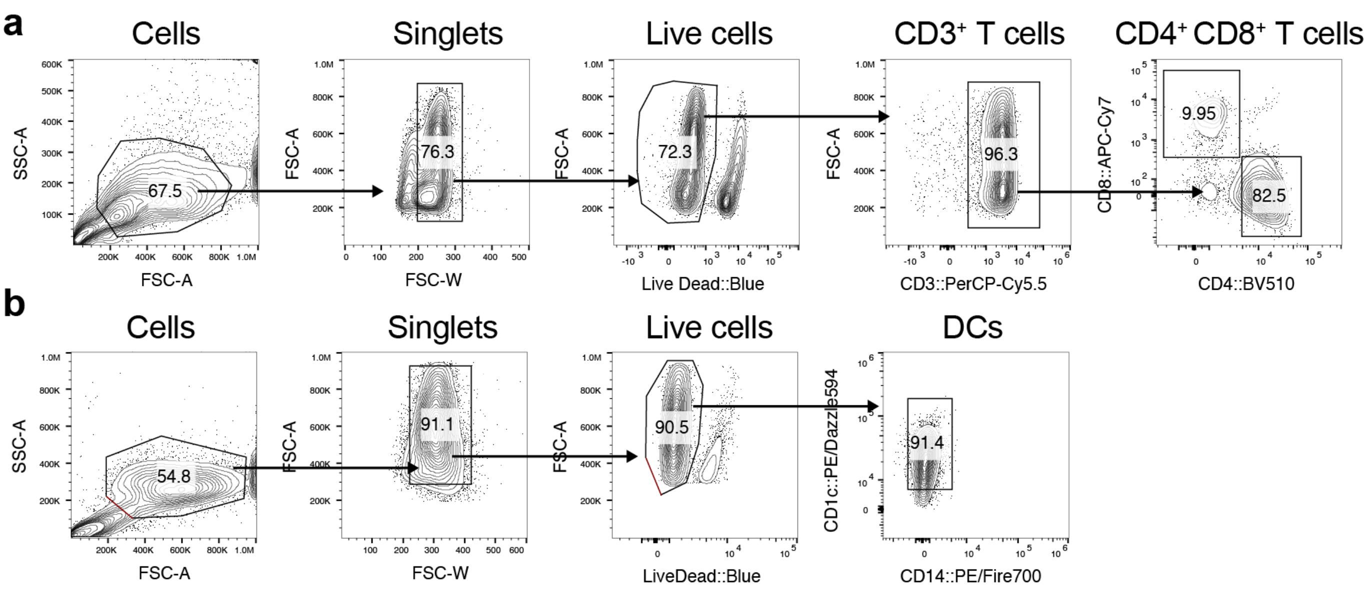


**Supplementary Fig. 13 | Representative flow cytometry gating strategy. Shown are the steps from the ungated population to CD4⁺ and CD8⁺ T cells (a) and to DCs (b).**

**Supplementary Movie 1 | DC-mediated collagen remodeling in 2 mg mL$^{-1}$ Fast R.**
DC nuclei (magenta) and actin (green) were labeled with SiR-DNA and SPY555-FastAct; collagen fibrils were labeled with tetrazine–iFluor 488. Time-lapse imaging was performed 2 h after encapsulation at 0.5 min per frame for 11 min. Scale bar, 10 μm; time, hh:mm.

**Supplementary Movie 2 | DC-mediated collagen remodeling in 2 mg mL$^{-1}$ Slow R.**
DC nuclei (magenta) and actin (green) were labeled with SiR-DNA and SPY555-FastAct; collagen fibrils were labeled with tetrazine–iFluor 488. Time-lapse imaging was performed 2 h after encapsulation at 0.5 min per frame for 11 min. Scale bar, 10 μm; time, hh:mm.

**Supplementary Movie 3 | p-a-blebbistatin-treated DC-mediated collagen remodeling in 2 mg mL$^{-1}$ Fast R.**
DC nuclei (magenta) and actin (green) were labeled with SiR-DNA and SPY555-FastAct, and collagen fibrils were labeled with tetrazine–iFluor 488. Time-lapse imaging was performed 2 h after encapsulation at 0.5 min per frame for 11 min. DCs were treated with 25 μM para-blebbistatin during incubation and throughout imaging. Scale bar, 10 μm; time, hh:mm.

**Supplementary Movie 4 | p-a-blebbistatin-treated DC-mediated collagen remodeling in 2 mg mL$^{-1}$ Slow R.**
DC nuclei (magenta) and actin (green) were labeled with SiR-DNA and SPY555-FastAct, and collagen fibrils were labeled with tetrazine–iFluor 488. Time-lapse imaging was performed 2 h after crosslinking at 0.5 min per frame for 11 min. DCs were treated with 25 μM para-blebbistatin during incubation and throughout imaging. Scale bar, 10 μm; time, hh:mm.

| ANTIBODIES | SOURCE | IDENTIFIER |
|---|---|---|
| Anti-human CD1c, PE/Dazzle594 | BioLegend | 331532 |
| Anti-human CD1c, PE/Cy5 | BioLegend | 331554 |
| Anti-human CD3, PerCP/Cy5.5 | BioLegend | 300430 |
| Anti-human CD4, BV510 | BioLegend | 317444 |
| Anti-human CD8, APC/Cy7 | BioLegend | 344714 |
| Anti-human CD14, PE/Fire700 | BioLegend | 399222 |
| Anti-human CD14, APC/Fire750 | BioLegend | 367120 |
| Anti-human CD25, PE/Cy5 | BioLegend | 302608 |
| Anti-human CD25, BV711 | BioLegend | 356138 |
| Anti-human CD40, APC/Cy7 | BioLegend | 313018 |
| Anti-human CD45RA, PE/Cy7 | BioLegend | 304126 |
| Anti-human CD80, PE/Cy5 | BioLegend | 305210 |
| Anti-human CD80, PE | BioLegend | 305208 |
| Anti-human CD83, APC | BioLegend | 305312 |
| Anti-human CD86, BV510 | BioLegend | 305432 |
| Anti-human CCR7, BV421 | BioLegend | 353208 |
| Anti-human CCR7, BV605 | BioLegend | 353224 |
| Anti-human PD-1, BV421 | BioLegend | 329920 |
| Anti-human PD-1, BV605 | BioLegend | 329924 |
| Anti-human PD-1, PE | BioLegend | 329906 |
| Anti-human PD-L1, BV785 | BioLegend | 329736 |
| Anti-human HLA-DR, APC | BioLegend | 307610 |
| Anti-human HLA-DR, FITC | BioLegend | 307604 |
| Anti-human HLA-ABC, FITC | BioLegend | 311404 |
| Anti-human OX-40, PE | BioLegend | 350004 |

**Supplementary Table 1 | Antibodies used for flow cytometry analysis.**
